# Immunometabolic role of metabolic enzyme PANK4 in regulation of TLR7/9-mediated innate immunity

**DOI:** 10.1101/2025.10.22.676442

**Authors:** Riya Chaudhary, Aparna Meher, Pratik Katekar, Drishiya Vats, Debasis Nayak, Himanshu Kumar

## Abstract

Innate immune responses are intimately linked to cellular metabolism, yet the molecular connections between metabolic reprogramming and antiviral defense remain incompletely defined. Here, we identify PANK4, an atypical member of pantothenate kinase family involved in coenzyme A (CoA) biosynthesis, as an unexpected regulator of RNA virus including influenza virus pathogenesis. Unlike other pantothenate kinases, PANK4 lacks canonical kinase activity but has been implicated in metabolic regulation. We show that influenza virus infection, in conjunction with pantothenic acid, induces PANK4 expression, which promotes viral replication by enhancing glucose uptake and glycolytic activity. Loss of PANK4 curtailed viral replication, reduced expression of glycolytic regulators, and heightened host antiviral defenses. Mechanistically, PANK4 interacts with UNC93B1 to suppress TLR7 and TLR9 mediated cytokine responses, thereby acting as a negative regulator of nucleic acid–sensing innate immune pathways in various cell-types. Furthermore, viral proteins NS1 and PB1 exploit PANK4 to amplify replication and immune evasion.

## INTRODUCTION

The innate immune system senses pathogens through pattern-recognition receptors (PRRs) that recognize conserved microbial signatures and initiate transcriptional programs essential for antiviral defense (Kumar *et al*, 2011). Endosomal Toll-like receptors (TLRs) and cytosolic RIG-I–like receptors (RLRs) sense viral RNA and trigger signalling cascades that culminate in type I interferon (IFN-I) and proinflammatory cytokine production. The magnitude and timing of these responses are tightly regulated to ensure efficient pathogen clearance while preventing immunopathology (Chaudhary *et al*, 2023).

Beyond canonical signalling components, cellular metabolism has emerged as a pivotal determinant in shaping innate immunity. Immune activation demands rapid bioenergetic and biosynthetic adaptation, and direct influence of metabolic enzymes on innate immune signalling pathways (Kelly & O’Neill, 2015). Viral pathogens frequently reprogram host metabolism to create a favourable environment for replication while evading immune surveillance. Increased glucose uptake and altered cofactor biosynthesis are common features of many viral infections, yet their precise impact on PRR-mediated antiviral immunity remains poorly defined (Bhatt *et al*, 2022; Goodwin *et al*, 2015).

Glycolysis initiates glucose catabolism, producing pyruvate that fuels acetyl-CoA synthesis, a central hub for generating essential biomolecules. Many viruses reprogram these pathways to support replication and drive pathogenesis (Thai *et al*, 2014). Synthesis of acetyl-CoA is tightly coupled to the pantothenic acid (PA)/vitamin B5 pathway, which drives the biosynthesis of CoA, a central cofactor in diverse metabolic and biosynthetic processes (Dibble *et al*, 2022).

PA imported into the cells primarily via the sodium-dependent multivitamin transporter SLC5A6 is subsequently phosphorylated by three enzymatically active and regulatory isoforms of pantothenate kinases (PANKs) in the first and rate-limiting step of CoA production (Yuan & Chen, 2025). Viruses exploit host metabolic intermediates and enzymes to remodel the metabolic landscape, enabling efficient resource utilization for replication while dampening antiviral innate immunity to facilitate immune evasion (Goodwin *et al*., 2015).

PANK4, a member of the pantothenate kinase family, diverges from its counterparts in lacking kinase activity and instead functions as a phosphatase, suggesting noncanonical regulatory roles (Dibble *et al*., 2022; Yao *et al*, 2019). The interplay between metabolites processed by PANK4, the resulting metabolic cues, and PANK4 itself, and how these collectively influence innate immune signalling pathways, remains unknown. Here, we identify metabolic–immune axis in which influenza A virus (IAV) infection enhances glucose uptake and increases PANK4 expression. We further show that PA, a CoA precursor, synergizes with IAV infection to markedly elevate PANK4 abundance, and that PA alone can upregulate PANK4 levels. Mechanistically, we demonstrate that PANK4 interacts with UNC93B1 and block-trafficking of TLRs essential for nucleic acid–sensing, including TLR7 and TLR9, to endolysosomal compartments for ligand recognition and signal initiation. Disruption of UNC93B1 function compromises antiviral responses and increases susceptibility to infection. Although UNC93B1 is subject to multilayered regulation, its connection to metabolic enzymes during viral infection has not been explored. Our findings uncover a direct link between viral metabolic reprogramming, and the control of PRR trafficking in antiviral immunity.

## RESULTS

### Identification of PANK4

To systematically identify host factors linking influenza virus replication with glucose metabolism, we integrated genome-wide screens of influenza virus host dependency factors (Konig *et al*, 2010) and kinase-focused IAV screens (Bakre *et al*, 2013) with a publicly available dataset of glucose-regulated genes in MDA-MB-231 cells (GSE225643) (Li *et al*, 2023). This analysis revealed 12 host factors that were both required for viral replication and upregulated in response to glucose. While most of these factors have been previously implicated in viral infection, PANK4 emerged as a previously uncharacterized regulator. PANK4 is highly conserved among mammalian species **(Fig. 1A and Fig. EV 1A–B, Appendix Table S 1)**. To evaluate whether glucose availability regulates PANK4, we analyzed normalized transcript counts (TPM) from the GSE225643 dataset. PANK4 expression was significantly higher in glucose-supplemented conditions compared to glucose starvation **(Fig. 1B)**. Analysis of PANK family members in mouse alveolar epithelial cells (mAECs) showed that PANK4 was selectively upregulated under glucose-supplemented compared to glucose-starved conditions **(Fig. 1C)**. We next examined PANK4 expression in A549 cells cultured under glucose-supplemented or glucose-starved conditions and observed a significant reduction in glucose-starved cells **(Fig. 1D)**. Immunofluorescence further revealed that PR8 infection induced PANK4 expression in a glucose-dependent manner **(Fig. 1E)**. Previously in one of screening that aimed at identifying host factors essential for viral replication (Bakre *et al*., 2013), PANK4 emerged as a prominent yet uncharacterized hit, prompting us to investigate its role in virus pathophysiology. PANK4 was upregulated in primary mouse cells, including mAECs following PR8 infection. In vivo, influenza virus infection also induced PANK4 expression in bronchoalveolar lavage (BAL) cells **(Fig. 1F–G and Fig. EV 2A, B)**. IAV is sensed by TLR7 and RIG-I, leading to IFN-I and inflammatory cytokine production to mount antiviral innate responses. We therefore evaluated whether PANK4 induction depends on IFN-I and inflammatory cytokine. PANK4 expression was not significantly induced in A549 cells treated with varying concentrations of IFN-β, although ISG15 was robustly induced in a dose-dependent manner. Consistently, IFN-β stimulation up to 48 hours failed to induce PANK4, whereas ISG15 remained strongly upregulated. To further assess IFN-I dependency, IRF3^-/-^ cells were examined; PANK4 expression was comparable in knockout (KO) and wild-type (WT) cells following IAV infection **(Fig. 1H-I and Fig. EV 2C)**.

**Figure 1.**
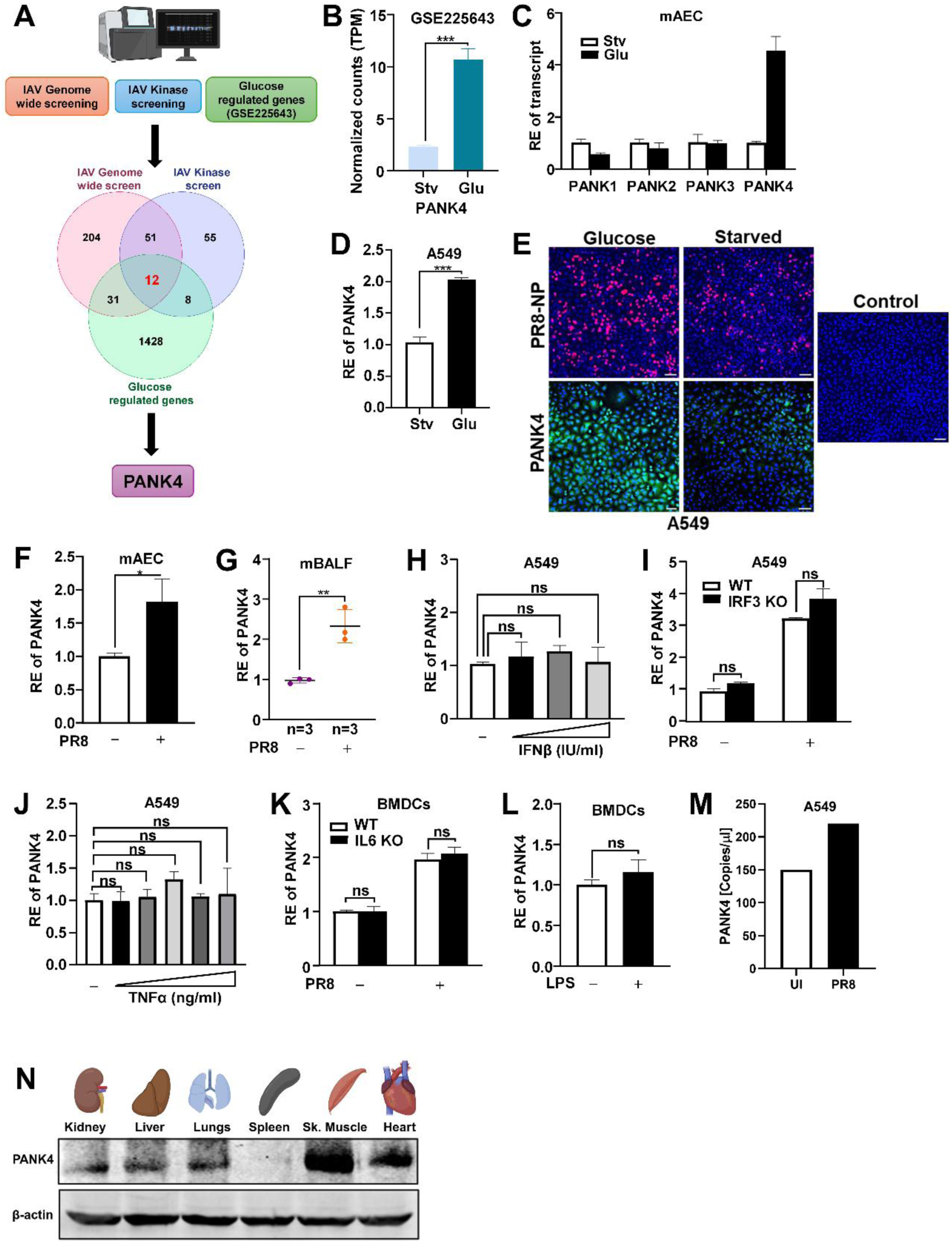
PANK4 is selectively induced by RNA virus infection and glucose, independent of type I IFN and proinflammatory cytokines. (A) Schematic representation of the integrative screening strategy combining genome-wide and kinase-focused influenza virus screens with a glucose-regulated gene dataset from MDA-MB-231 cells. (B) TPM values for PANK4 expression in MDA-MB-231 cells cultured under glucose-starved (Stv) or glucose-supplemented (Glu) conditions from GSE225643. (C) mAECs were cultured under glucose-supplemented versus glucose-starved (12 h) conditions and RNA level of PANK family members were analysed. (D) A549 cells were cultured under glucose-supplemented or glucose-starved conditions (12h) and PANK4 expression was measured. (E) A549 cells under glucose-supplemented or glucose-starved (12 h) conditions were infected with PR8 (MOI 1) for 12 h. PR8-NP and PANK4 were analysed by immunofluorescence microscopy using indicated antibodies. (F–G) Ex vivo, mAECs were isolated and infected with PR8 (MOI 1) for 24 h. In vivo, BAL fluid (G) (PFU 100, dpi 3) was collected, after which the RNA level of PANK4 was measured. (H) A549 cells were treated with increasing concentrations of IFN-β (100, 200, and 500 IU/ml) for 24 h and PANK4 RNA level were analysed. (I) WT and IRF3^-/-^ A549 cells with or without PR8 (MOI 1) 24 h post infection PANK4 transcript level was measured. (J) A549 cells stimulated with TNF-α (10– 100 ng/ml) for 12 h and analysed for PANK4 transcript level. (K) WT and IL6^-/-^ BMDCs infected with PR8 (MOI 1) or without infection. (L) BMDCs stimulated with LPS (100 ng/ml) for 24 h and PANK4 expression was measured. (M) A549 cells were infected with PR8 (MOI 1) and absolute PANK4 transcript copy number were determined by ddPCR. (N) Immunoblot representing the tissue distribution of PANK4 in mice. C-D and F-L were analysed by RT-PCR. Data are presented as the mean ± SEM from triplicate samples of a single experiment and representative of three to five independent experiments. *** p <0.001, ** p <0.01, * p<0.05 and ns is non-significant by two-tailed unpaired Student’s t-tests.

Next, we tested whether inflammatory cytokines regulate PANK4. Treatment of cells with varying TNF-α concentrations failed to induce PANK4 but strongly upregulated IL-6. Moreover, IL-6^-/-^ and WT BMDCs exhibited comparable PANK4 induction. Notably, LPS stimulation of BMDCs did not elevate PANK4 expression. Moreover, PANK4 transcript copy number was significantly increased in virus-infected cells **(Fig. 1J-M and Fig. EV 2D-F)**.

Finally, we profiled tissue distribution of PANK4 in mice. PANK4 was ubiquitously expressed, with highest levels in skeletal muscle and heart, and similarly expressed in human tissues **(Fig. 1N and Fig. EV 2G)**.

Collectively these findings indicating, IAV infection specifically induces PANK4 in IFN-I and inflammatory cytokines independent manner.

### Immunometabolic role of PANK4 in RNA virus infection

Previously, it has been shown that the viral replication is suppressed under glucose deprivation (Bhatt *et al*., 2022), we examined viral load following infection under glucose-supplemented and glucose starved conditions and found that the presence of glucose significantly enhanced viral replication **(Fig. 2A)**. In a recent report, PANK4 identified as a metabolic regulator that enhances glucose uptake in skeletal muscles (Miranda-Cervantes *et al*, 2025). Since viruses are known to manipulate host glucose metabolism to facilitate viral replication, we investigated PANK4 role in virus-induced metabolic reprogramming (Bhatt *et al*., 2022). Notably, among all isoforms, only PANK4 showed consistent induction upon PR8 infection in both mAECs and HEK293T cells under glucose-supplemented conditions **(Fig. EV 3A–B)**, suggesting that PANK4 uniquely responds to both viral infection and glucose availability.

**Figure 2.**
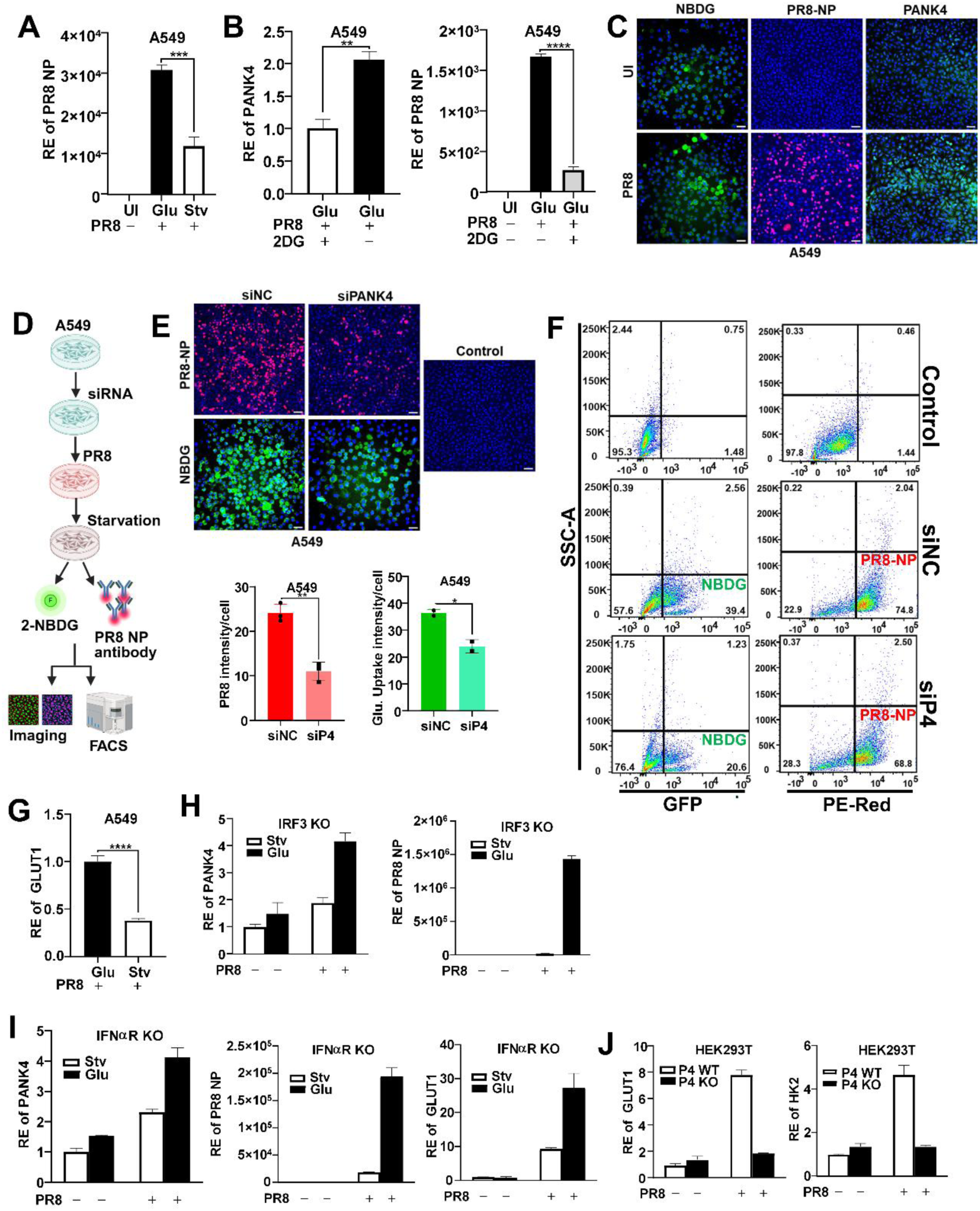
PANK4 mediates virus-induced glycolytic reprogramming to support viral replication. (A) A549 cells were cultured under glucose-supplemented versus glucose-starved (12 h) conditions and RNA level of PR8-NP were analysed. (B) A549 cells were infected with PR8 (MOI 1) for 12 h and subjected to 2-DG (5 mM) for 12 h treatment. The RNA level of PANK4 and PR8-NP were analysed. (C) A549 cells were infected with PR8 (MOI 1, 24 h) and incubated with 2-NBDG (200 µM, 20 min, 37 °C). 2-NBDG, PR8-NP and PANK4 were analysed by Immunofluorescence microscopy. (D–F) A549 cells were transfected with siNC (control) or siPANK4 (30 nM) and infected with PR8 (MOI 1, 12 h) and subjected to starvation (6 h). Glucose uptake (2-NBDG) and viral load (PR8-NP antibody) were measured by (E) immunofluorescence microscopy and (F) flow cytometry. Plots show 2-NBDG fluorescence and NP staining, with quadrants representing NP and NBDG populations; numbers denote percentage of cells. (G) A549 cells were cultured in glucose-supplemented versus glucose-starved (12 h) and infected with PR8 (MOI 1, 12 h), RNA level of GLUT1 were measured. (H-I) IRF3⁻/⁻ A549 cells and IFNαR⁻/⁻ BMDCs were cultured in glucose-supplemented versus glucose-starved (12 h) or infected with PR8 (MOI 1, 12 h). Transcript level of PANK4 and PR8-NP were measured in IRF3⁻/⁻ A549 cells and transcript level of PANK4, PR8-NP and GLUT1 were measured. (J) WT and PANK4⁻/⁻ HEK293T (PANK4⁻/⁻) cells were infected with PR8 (MOI 1, 24 h) after that RNA level of GLUT1 or HK2 were analysed. A, B, G, H, I and J were analysed by RT-PCR. Data are presented as the mean ± SEM from triplicate samples of a single experiment and representative of three to five independent experiments. **** p<0.0001, *** p<0.001, ** p<0.01, * p<0.05 by two-tailed unpaired Student’s t-tests.

To further dissect the relationship between glycolysis and PANK4 during viral infection, we treated glucose-supplemented, PR8-infected A549 cells with 2-deoxy-D-glucose (2DG), well-characterized competitive inhibitor of glycolysis. 2DG is a glucose analogue that is taken up by glucose transporters such as glucose transporter 1 (GLUT1) and phosphorylated by hexokinase (HK2) to form 2DG-6-phosphate, which cannot be further metabolized through the glycolytic pathway. This accumulation inhibits downstream glycolytic enzymes and disrupts glycolytic flux, thereby mimicking a glucose-deprivation state despite the presence of extracellular glucose(Pajak *et al*, 2019). Treatment with 2DG led to a marked suppression of PANK4 expression, suggesting that active glycolytic metabolism is essential for the virus-induced upregulation of PANK4 **(Fig. 2B)**. These findings highlight the dependence of PANK4 expression on host glycolytic activity during infection. To assess whether influenza virus infection enhances glucose uptake and whether this correlates with PANK4 expression, we monitored the uptake of the fluorescent glucose analogue 2-NBDG under glucose-starved conditions during infection. PR8-infected cells exhibited marked increase in 2-NBDG uptake, which was accompanied by elevated PANK4 expression **(Fig. 2C)**.

To further investigate PANK4 role in glucose uptake, we first silenced PANK4 expression, followed by PR8 infection in glucose supplemented or starvation condition. Under these conditions, loss of PANK4 resulted in marked reduction in both viral load and glucose uptake, as determined by fluorescence microscopy and flow cytometry, in both infected and uninfected cells **(Fig. 2D–F and Fig. EV 3C–D)**. Conversely, overexpression of PANK4 enhanced glucose uptake **(Fig. EV 3E)**. These findings indicate that PANK4 promotes glucose uptake and facilitate viral replication by modulating host glycolytic pathways. To dissect the mechanism underlying virus-induced alterations in glucose metabolism, we examined the expression of GLUT1, key regulator of glycolytic flux. Elevated expression of GLUT1 upon PR8 infection under glucose-supplemented conditions suggests an active metabolic reprogramming toward glycolysis to support viral replication **(Fig. 2G)**.

To examine whether this metabolic regulation was dependent on IFN-I signalling, we used IRF3^-/-^ A549 cells and IFNαR^-/-^ BMDCs, as IRF3 is a key transcription factor for IFN-I production, while IFNαR is essential for IFN receptor–mediated signalling. PANK4 induction was solely depends on availability of glucose and virus infection in IFN-independent manner and the expression of GLUT1 linked to glucose rather than IFNs **(Fig. 2H–I)**.

Finally, PR8 infection led to significant upregulation of GLUT1 and HK2 in WT but not in PANK4^-/-^ cells, with the most robust induction observed in glucose-fed WT cells **(Fig. 2J and Fig. EV 3F)**. Together, these results identify PANK4 as a critical mediator of virus-induced glycolytic reprogramming and viral replication, functioning independently of IFN-I signalling axis.

### PA induces PANK4 and supports viral pathogenesis in vivo

Glucose metabolism is tightly linked to CoA biosynthesis, as the intermediates generated from glycolysis feed into various CoA-dependent pathways. CoA is synthesized from PA, which is taken up by cells via specific transporter (SLC5A6) and phosphorylated by PANK1, 2, & 3 whereas PANK4 is a regulatory isoform and inhibit other PANKs (Yao *et al*., 2019). So, we focused on how this metabolic shift may influence PA utilization and PANK4 function. Having shown that PANK4 promotes virus-induced glucose uptake and supports viral replication through metabolic reprogramming, we next investigated whether this response could be modulated by PA **(Fig. 3A)**. We first examined PANK4 expression over time following PA treatment by immunoblot and observed substantial increase up to 12 hours, with expression levels remaining stable up to 36 hours, suggesting that PA induces a sustained upregulation of PANK4 **(Fig. 3B)**. A similar pattern of PANK4 upregulation was observed with increasing concentrations of PA followed by PR8 infection, indicating PA enhances the PANK4 expression **(Fig. EV 4A-B).** In A549 cells, immunofluorescence revealed increased PANK4 expression upon PA treatment **(Fig. 3C)**. Consistent with this, we observed similar upregulation of PANK4 in multiple primary human and mouse cell types including PBMCs, BMDCs, mouse lung tissue, and human cell lines HEK293T, and THP-1 cells—by immunoblot, indicating a broadly conserved response to PA across both human and mouse systems **(Fig. 3D-E)**. Next, we observed PA treatment further enhanced PANK4 expression in the context of PR8 infection in PBMCs and A549 cells, which was accompanied by increase in viral load, suggesting PA may facilitate viral replication through PANK4 induction. Additionally, SLC5A6 was found upregulated in both PBMCs and A549 cells following PA treatment **(Fig. 3F-G)**. We observed significantly enhanced SLC5A6 expression in mouse lung tissues upon infection **(Fig. 3H).** To test PA utilization from the culture medium, we also examined SLC5A6 expression in A549 cells overexpressing PANK4. PANK4 overexpression led to marked increase in SLC5A6 levels, suggesting enhanced PA uptake. Conversely, PANK4^-/-^ cells resulted in significantly reduced SLC5A6 expression, indicating a potential relation between PANK4 activity and PA transport **(Fig. 3I-J)**. To validate in vitro findings in physiological setting, we employed an in vivo mouse model to explore the relationship between PANK4 and PA during virus infection and selected outbred CD1 mice. Mice were administered with PA and infected as shown in schematic **(Fig. 3K)**. We observed a significant increase in viral load in the PA-treated and virus-infected group compared to the group infected with virus alone, indicating that PA supplementation enhances viral replication in vivo **(Fig. 3L)**. Next, we examined PANK4 expression in lung tissues and found that it was elevated in both PA-treated and virus-infected groups individually, with the highest expression observed in the PA plus virus group. This suggests synergistic effect of PA and viral infection on PANK4 induction **(Fig. 3M)**. In parallel, we tested SLC5A6 expression **(Fig. 3N)**. These findings were further validated by immunoblot analysis, confirming the protein-level changes in both PANK4 and viral load **(Fig. 3O)**. To examine the impact of virus and PA, we monitored body weight daily as an indicator of pathogenesis **(Appendix Table S 2)**. Mice receiving PA plus virus treatment showed significant reduction in body weight compared to those infected with virus alone, suggesting that PA supplementation exacerbates pathogenesis during virus infection **(Fig. 3P and Fig. EV 4C)**. In summary, these findings demonstrate that PA enhances PANK4 expression and promotes influenza virus replication, likely by modulating host glucose and CoA metabolism.

**Figure 3.**
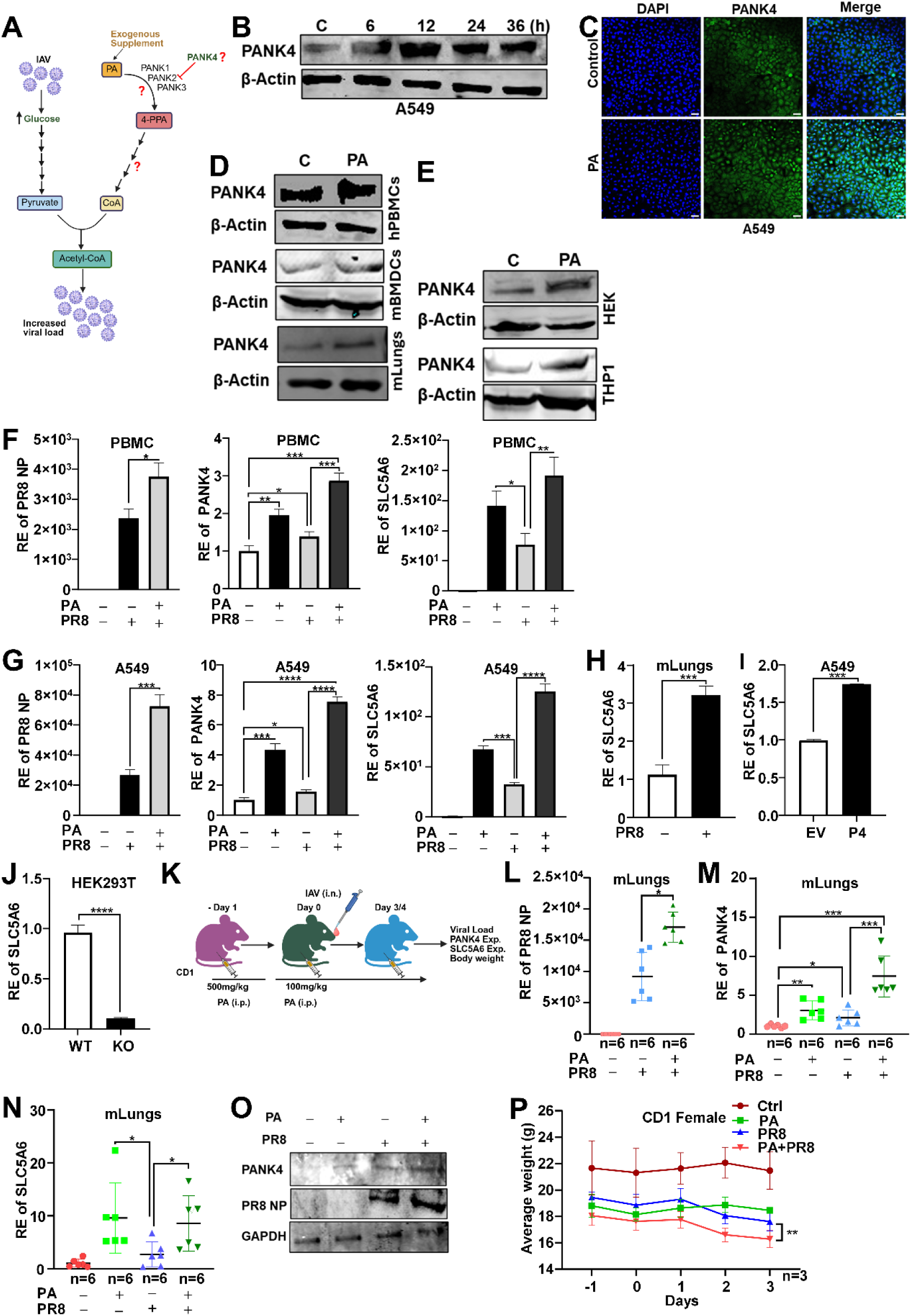
PA enhances PANK4 expression and promotes viral replication in vitro and in vivo. (A) Schematic representation of glucose and PA metabolism and CoA biosynthesis highlighting the involvement of PANK4. (B) A549 cells treated with PA (100 µM) and PANK4 expression analysed by immunoblot over indicated time-course. (C) A549 cells treated with PA (100 µM, 24 h) and PANK4 expression analysed by immunofluorescence microscopy (scale bar, 100 µm). (D) Human (h) PBMCs, mouse (m) BMDCs, and mouse lung tissue, mouse were given PA 500mg/kg, i.p. for 24 h (E) human cell lines (HEK293T or THP1) treated with PA (100 µM, 24 h) and PANK4 expression was analysed by immunoblot. (F, G) PBMCs or A549 cells treated with PA (100 µM, 24 h) and RNA level of SLC5A6, PR8-NP, and PANK4 were analysed. (H-I) C57BL/6 Mice were infected with PR8 (PFU 100, dpi 3) lung tissue were collected; A549 cells were transfected with PANK4 plasmid and infected with PR8 (MOI 1, 24 h). RNA level of SLC5A6 were analysed. (J) WT or PANK4⁻/⁻ cells were cultured and RNA level of SLC5A6 were analysed. (K) Schematic of in vivo infection model in CD1 mice: prior to intranasal PR8 infection (PFU 100), i.p.of PA (500mg/kg) was given and subsequently maintained with 100 mg/kg as indicated, and lung tissue collection at dpi 3. Transcript levels of PR8 NP (L), PANK4 (M), and SLC5A6 (N) were analysed. (O) Protein levels of PR8 NP and PANK4 analysed by immunoblot. (P) Female mice body weight changes measured over the indicated time course. F-J and L-N were analysed by RT-PCR. Data are presented as the mean ± SEM from triplicate samples of a single experiment and representative of two to three independent experiments. **** p<0.0001, *** p<0.001, ** p<0.01, * p<0.05 by two-tailed unpaired Student’s t-tests or two-way repeated-measures ANOVA test.

### PANK4 enhances viral replication through suppression of innate antiviral responses

Virus infection, glucose and PA induced PANK4 expression, to examine how PANK4 modulates viral infection, A549 cells were infected with increasing MOIs of IAV for 24 h. PANK4 expression and viral load were measured by immunoblotting using PANK4 and IAV-specific antibodies, and by RT-PCR at transcript level. Interestingly, PANK4 expression increased proportionally with viral load **(Fig. 4A and Fig. EV 5A)**. Similar, results were observed in RAW264.7 cells, mouse BMDCs as analysed by immunoblot **(Fig. 4B-C and Fig. EV 5B).** Moreover, PBMCs and THP-1 cells also show similar trend following IAV infection **(Fig. 4D-E)**. Furthermore, A549 cells infected with NDV also show similar results **(Fig. EV 5C)**.

**Figure 4.**
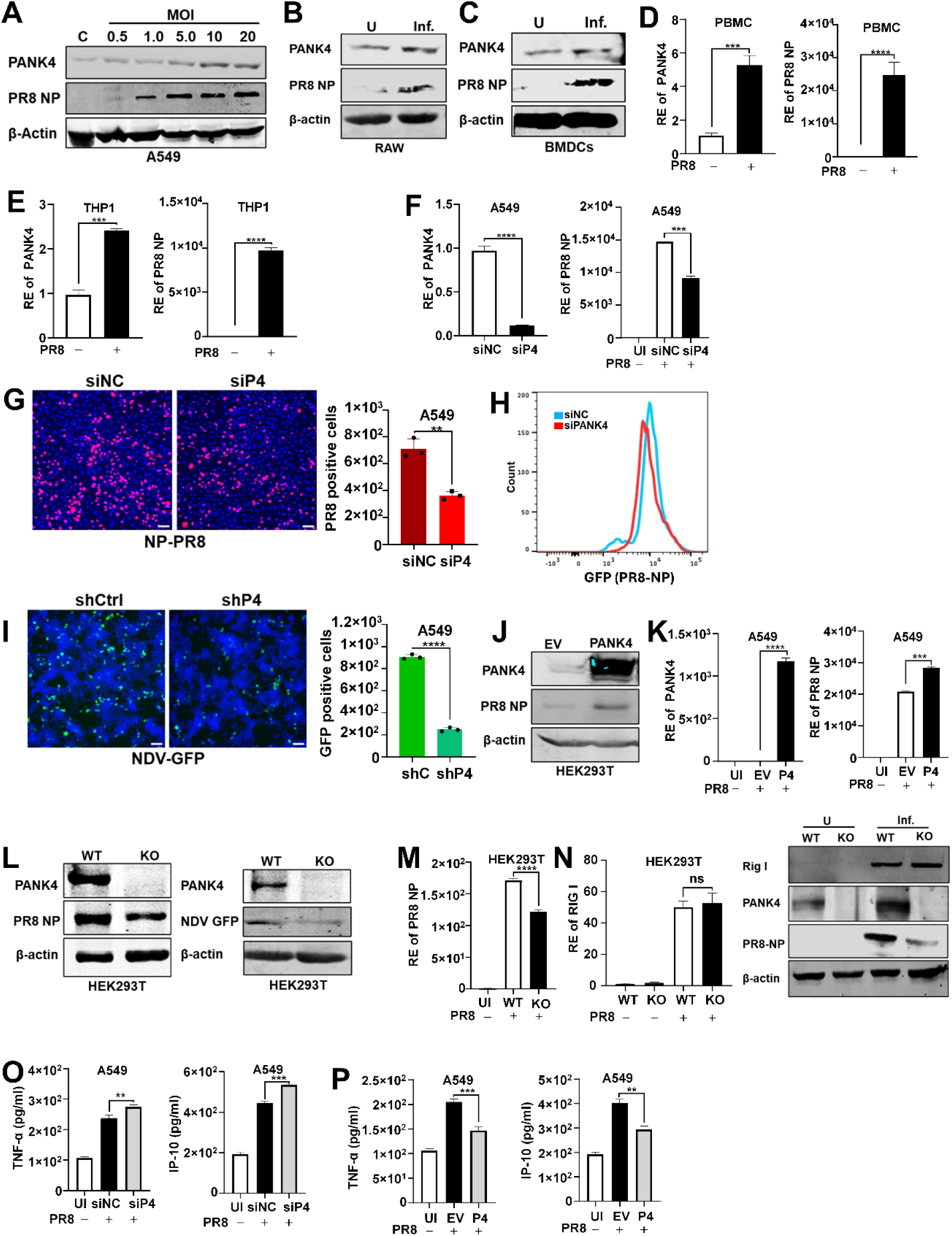
PANK4 promotes RNA virus replication and regulates host cytokine responses. (A) A549 cells were infected with PR8 at the indicated MOIs for 24 h and PANK4 and PR8 NP expression analysed by immunoblot. (B) RAW264.7 cells (RAW) and (C) BMDCs infected with PR8 (MOI 1, 24 h) and protein level of PANK4 and PR8-NP were analysed by immunoblot. (D) human PBMCs, and (E) THP1 cells were infected with PR8 (MOI 1, 24 h) and RNA level of PANK4 were analysed. (F–H) A549 cells were transfected with siPANK4 and infected with PR8 (MOI 1, 24 h) and PR8-NP expression analysed by (F) RT-PCR, (G) immunofluorescence microscopy, and (H) flow cytometry. (I) A549 cells were transfected with shRNA of PANK4 and infected with NDV-GFP (MOI 0.5, 24 h) and NDV–GFP level were analysed by immunofluorescence. (J-K) HEK293T and A549 cells were transfected with construct encoding PANK4 and infected with PR8 (MOI 1, 24 h). PR8-NP and PANK4 protein level were measured by (L) immunoblot and (M) RT-PCR. (N) WT or PANK4⁻/⁻ cells were infected with PR8 (MOI 1, 24 h) or NDV-GFP (MOI 0.5, 24 h) and NP or NDV–GFP (α-GFP) expression were analysed by immunoblot. (M) WT or PANK4⁻/⁻ cells were infected with PR8 (MOI 1, 24 h) and PR8-NP were analysed. (N) WT and PANK4⁻/⁻ cells were infected with PR8 (MOI 1, 24 h) and RIG-I expression were analysed by RT-PCR and immunoblot. (O-P) A549 cells were cultured and transfected with siPANK4 (O) or PANK4 overexpression plasmid (P) and infected with PR8 (MOI 1). After 24 h, the concentration of the TNF-α and IP-10 in the culture supernatant was measured by ELISA. D-F and M-N were analysed by RT-PCR. Data are presented as the mean ± SEM from triplicate samples of a single experiment and representative of three to five independent experiments. **** p<0.0001, *** p<0.001, ** p<0.01, ns is non-significant by two-tailed unpaired Student’s t-tests.

To examine the effect of PANK4 on virus replication, PANK4 knockdown (KD) was performed by siRNA in A549 cells followed by IAV infection, PANK4 KD cells showed a marked reduction in influenza viral load compared to the control as tested by RT–PCR, immunofluorescence, and flow cytometry; a similar decrease in NDV replication was also observed by immunofluorescence and RT–PCR following PANK4 silencing following NDV infection **(Fig. 4F–I and Fig. EV 5D)**. In contrast, PANK4 overexpression in HEK293T and A549 cells enhanced viral replication **(Fig. 4J–K and Fig. EV 5E–F)**. Furthermore, these results were confirmed using PANK4^-/-^ cells which show significant reduction of viral load upon IAV or NDV infection as shown by immunoblot, using specific antibody and examined by RT-PCR. However, expression of RIG-I was comparable in WT and PANK4^-/-^ cells as tested for RIG-I by RT-PCR and immunoblot **(Fig. 4M–N)**. Notably, positive sense RNA virus, DENV infection demonstrated significant viral reduction in PANK4^-/-^ cells compared to WT cells. Moreover, DENV replication enhanced in PANK4 overexpressed cells compared to empty vector introduced into liver cell line, HepG2 **(Fig. EV 5H–I)**.

Finally, innate immune responses in terms of TNF-α and IP-10 production were analysed at protein level in culture supernatant by ELISA after PANK4 KD or overexpression followed by the IAV infection. PANK4 KD show significantly enhanced production of TNF-α and IP-10 whereas PANK4 overexpression significantly reduced these cytokines **(Fig. 4O–P)**.

Overall, these results suggest that RNA virus infection induces PANK4 and support viral replication in various cell-types through reduction of antiviral innate immune responses.

### PANK4 interact with UNC93B1 and modulate TLR7/9 signalling pathway

Recent studies have highlighted that metabolic enzymes can regulate immune signalling independently of classical antiviral roles. Given that PANK4 is upregulated during influenza virus infection and is known to modulate cellular metabolism, we hypothesized that PANK4 may influence innate immune signalling by interacting with key host factors. Since innate immunity relies on tightly regulated protein-protein interactions for the sensing of viral components and initiation of downstream signalling cascades, we sought to identify PANK4-associated proteins that could reveal novel regulatory roles in this context. To investigate the potential involvement of PANK4 in innate immune signalling, we first performed an in-silico screen to identify its putative interacting partners. Using BioGRID, we identified 84 PANK4-interacting proteins (Oughtred *et al*, 2021). Cross-referencing with InnateDB revealed 14 overlapping genes linked to innate immune pathways, suggesting a potential role for PANK4 in innate immune regulation **(Fig. 5A, Appendix Table S 3)**. Among the 14 candidates, we focused on UNC93B1, a key regulator of the TLR7 and TLR9 signalling pathway and a critical mediator of antiviral innate immune responses. To validate the interaction between PANK4 and UNC93B1, we performed Flag-antibody-based co-immunoprecipitation assays under both uninfected and virus-infected conditions. Endogenous PANK4 was found to interact more robustly with UNC93B1 and TLR7 during influenza virus infection in A549 cells **(Fig. 5B)**. To further confirm this interaction, we overexpressed FLAG-tagged UNC93B1 or RIG-I along with MYC-tagged PANK4 and observed a specific interaction between PANK4 and UNC93B1 but not with RIG-I **(Fig. EV 6A)**. Additionally, immunofluorescence revealed co-localization of PANK4 and UNC93B1, with a high Pearson’s correlation coefficient, supporting their spatial proximity during infection **(Fig. 5C and Fig. EV 6B)**. Next, we examined the endogenous expression of UNC93B1 and TLR7 in THP-1 cells following PANK4 silencing and IAV infection. Upon PANK4 knockdown, we observed increased expression of UNC93B1 and TLR7 at both mRNA and protein levels, as confirmed by RT-PCR and immunoblot analysis **(Fig. 5D-E)**. Conversely, overexpression of PANK4 in THP-1 cells led to reduced expression of UNC93B1 and TLR7 at mRNA level **(Fig. EV 6C)**. Notably, the siRNA sequence targeting PANK4 is conserved between human and mouse **(Fig. EV 6D)**. PANK4 knockdown in RAW cells increased UNC93B1 and TLR7 expression at both mRNA and protein levels, with similar protein-level effects observed in BMDCs **(Fig. 5F-H)**. Next, using PANK4^-/-^, we observed consistent results, elevated UNC93B1 and TLR7 expression confirmed by immunoblot, RT-PCR, and immunofluorescence for UNC93B1 **(Fig. 5I-K)**. Moreover, PANK4 was overexpressed in PBMCs, which resulted in reduced expression of UNC93B1, TLR7 and enhanced viral load **(Fig. EV 6E)**. CpG ODN stimulation increased TLR9 in RAW cells, and in A549 cells it induced UNC93B1, TLR9, and IL-6 without affecting PANK4 expression (**Fig. EV 6F−G)**. Upon PANK4 knockdown in BMDCs and RAW cells stimulated with CpG ODN, we observed increased expression of both UNC93B1 and TLR9 at the mRNA level, as confirmed by RT-PCR. Moreover, it enhances IL-6 expression in RAW cells **(Fig. 5L–N)**. Consistently, increased expression of UNC93B1 and TLR9 upon PANK4^-/-^ was also observed by RT-PCR **(Fig. EV 6H)**. Together, these findings demonstrate that PANK4 interacts with UNC93B1 and negatively regulates TLR7 and TLR9 signalling pathways, revealing previously unrecognized role of PANK4 in modulating innate immune responses during viral infection.

**Figure 5.**
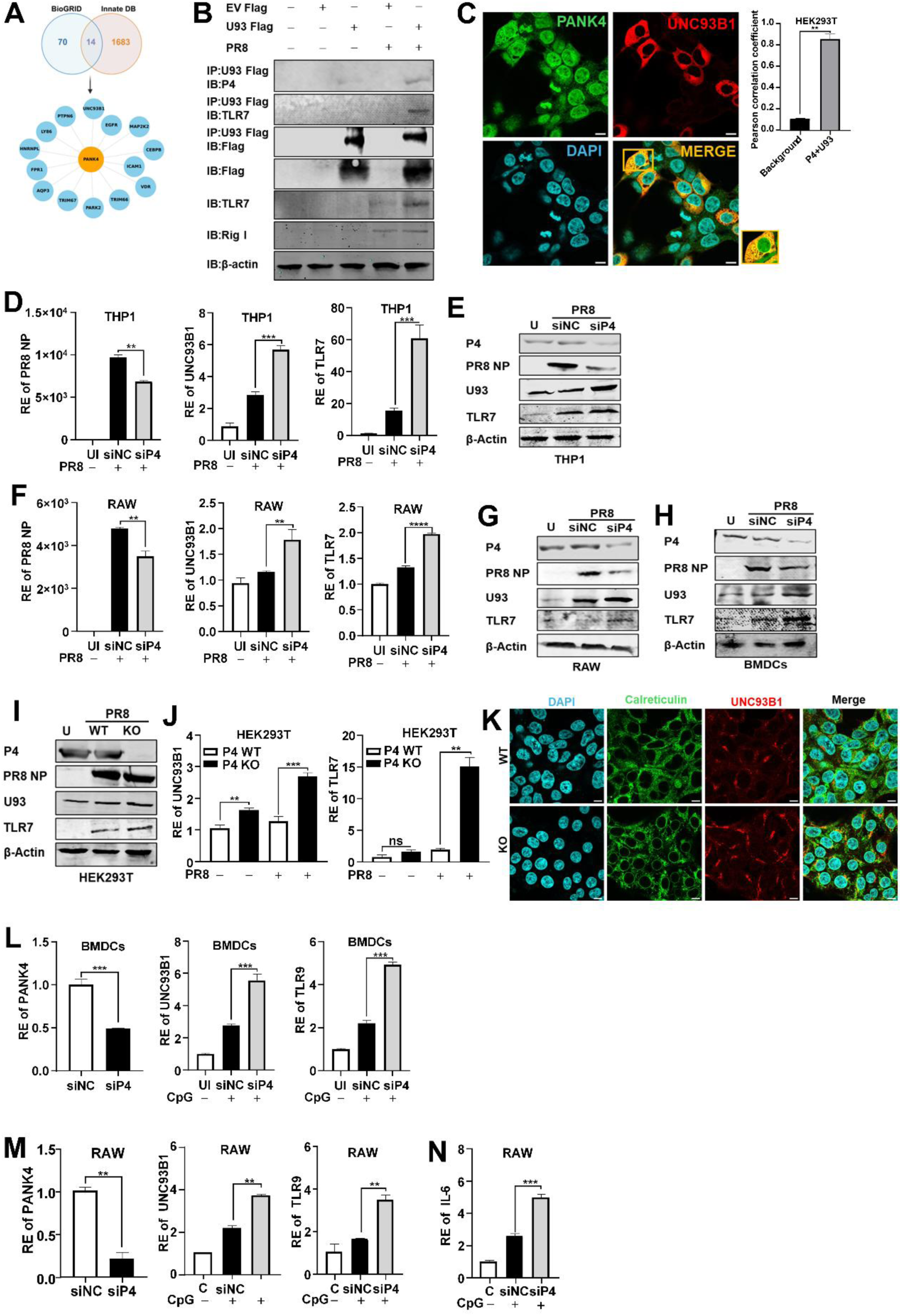
PANK4 interacts with UNC93B1 and negatively regulates TLR7 and TLR9 signalling. (A) Venn diagram showing overlap between PANK4 interactors identified in BioGRID and innate immune genes curated in InnateDB. (B) A549 cells transfected with Flag-tagged UNC93B1 (U93) subsequently infected with PR8 (MOI 1, 24 h) and proceeded for co-immunoprecipitation of endogenous PANK4 (P4) with UNC93B1–Flag; TLR7 included as positive control. (C) HEK293T cells were infected with PR8 (MOI 1, 24 h) and co-localization of PANK4 and UNC93B1 analysed by immunofluorescence microscopy (scale bars, 10 μm) and quantified by Pearson’s correlation coefficient. (D, E) THP1 cells were electroporated with siPANK4 and infected with PR8 (MOI 1, 24 h) and PR8-NP, UNC93B1 and TLR7 expression analysed by (D) RT-PCR and (E) immunoblot. (F and G) RAW cells and (H) BMDCs were transfected with siPANK4 and infected with PR8 (MOI 1, 24 h) and PR8-NP, UNC93B1 and TLR7 expression analysed by (F) RT-PCR and (G-H) immunoblot. (I–J) WT or PANK4⁻/⁻ cells were cultured or infected with PR8 (MOI 1, 24 h) and PR8-NP, UNC93B1 and TLR7 expression analysed by (I) immunoblot, UNC93B1 and TLR7 expression analysed by (J) RT-PCR. (K) WT or PANK4⁻/⁻ cells infected with PR8 (MOI 1, 24 h) and UNC93B1 expression were analysed by immunofluorescence microscopy (scale bars, 10 μm). (L–N) BMDCs and RAW cells were transfected with siPANK4 and stimulated with CpG ODN (5 µg/mL, 24 h) and expression of UNC93B1, TLR9, and IL-6 were analysed. L-N were analysed by RT-PCR. Data are presented as the mean ± SEM from triplicate samples of a single experiment and representative of three independent experiments. **** p<0.0001, *** p<0.001, ** p<0.01, * p<0.05 by two-tailed unpaired Student’s t-tests.

### PANK4 full length protein is essential for function

PANK4 interacts with UNC93B1, we sought to determine whether this interaction requires the full-length protein or can be mediated by individual domains. Notably, the PANK4 consist of N-terminal kinase which is nonfunctional and C-terminal phosphatase domain. To address this, we generated N-terminal and C-terminal truncated mutants lacking kinase (ΔC) and phosphatase (ΔN) domain of PANK4, respectively **(Fig. 6A)** and assessed their ability to bind UNC93B1. To this end, co-immunoprecipitation was performed using PANK4 and its mutants with UNC93B1, the analysis showed that both the ΔN and ΔC domains retained the capacity to interact with UNC93B1 **(Fig. 6B)**. However, overexpression of full-length PANK4 enhanced viral load, whereas neither the ΔN nor ΔC domain alone was able to do so in A549 cells as tested by RT-PCR **(Fig. 6C)**. Similar results were obtained by immunofluorescence and immunoblot which likewise showed increased viral load only in the presence of full-length PANK4 **(Fig. 6D, E)** indicating that the intact protein is required for regulation of virus.

**Figure 6.**
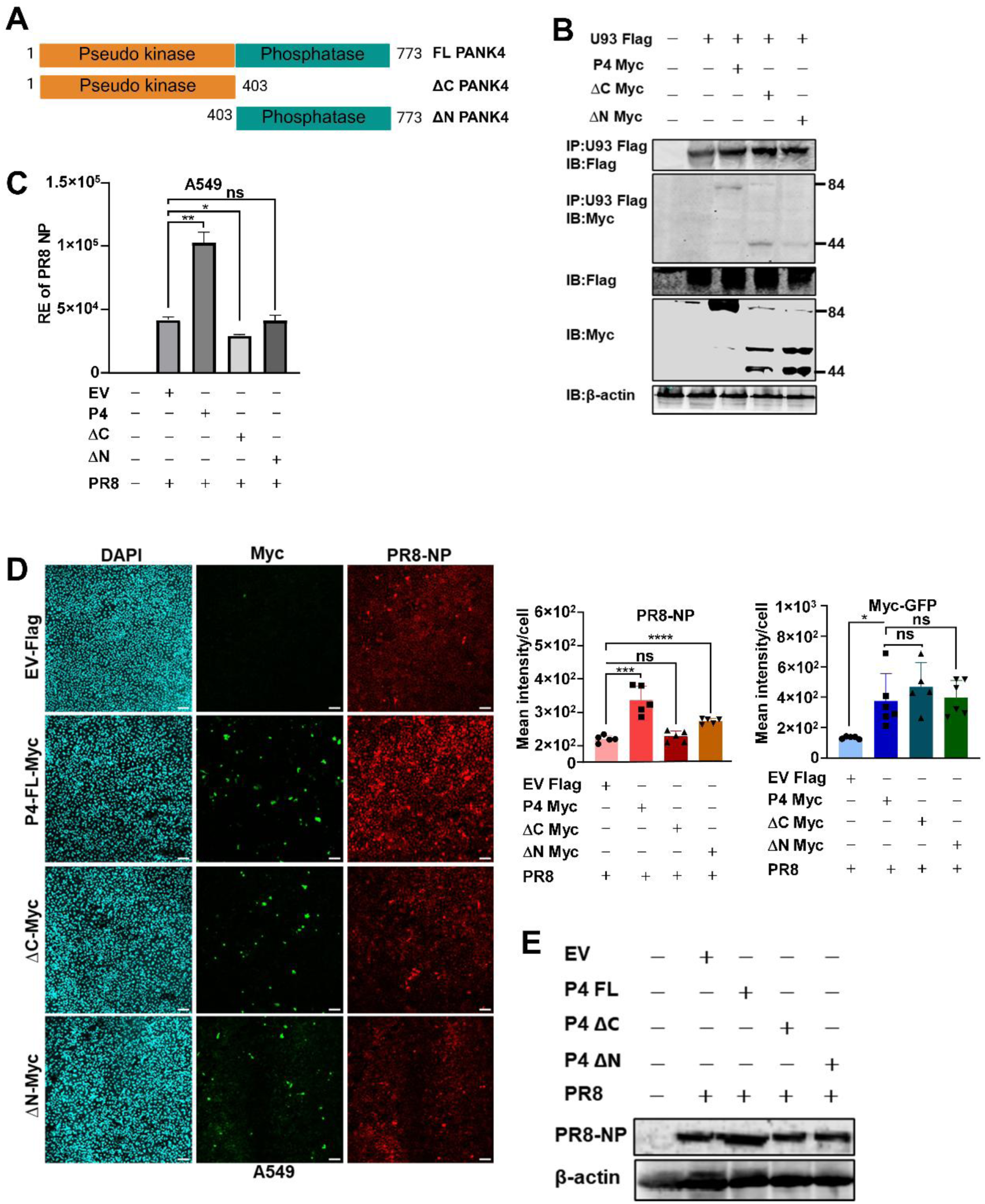
Full-length PANK4 is required for functional activity during viral infection. (A) Schematic representation of full-length PANK4 and truncated mutants lacking either the C-terminal phosphatase domain (ΔC PANK4) or the N-terminal kinase-like domain (ΔN PANK4). (B) HEK293T cells were transfected with overexpressing full-length PANK4, ΔN, or ΔC mutant (MYC-tagged) plasmids and subjected for co-immunoprecipitation with FLAG-tagged UNC93B1 and analysed by immunoblot (C) A549 cells were transfected with overexpressing full-length PANK4, ΔN, or ΔC mutant plasmids and infected with PR8 (MOI 1, 24 h). PR8-NP expression were analysed by RT-PCR (D−E) A549 cells were transfected with overexpressing full-length PANK4 or truncated mutant plasmids. PR8-NP and PANK4 expression were analysed by immunofluorescence microscopy (scale bar, 100 µm) and immunoblot; mean fluorescence intensity per cell quantified using QuPath-0.4.3 from five independent replicates. Data are presented as the mean ± SEM from triplicate samples of a single experiment and representative of three independent experiments. ** p<0.01, * p<0.05, ns is non-significant by two-tailed unpaired Student’s t-tests.

### PANK4 interact with IAV NS1 and PB1 and enhance viral replication

Viral proteins often exploit host factors to promote replication and suppress immune defences. Since PANK4 has emerged as a regulator in viral infection, we examined its potential interactions with IAV proteins. In silico prediction using the Human Virus Interaction Database (HVIDB) (Yang *et al*, 2021) indicated potential interactions of PANK4 with influenza proteins PB1 and NS1 **(Fig. 7A)**. Co-immunoprecipitation screening of PANK4 with four influenza proteins (PB1, NS1, M2, and NP) revealed specific interactions with PB1 and NS1 **(Fig. EV 7A)**. These results were corroborated by focused co-immunoprecipitation assays, which confirmed that PANK4 specifically associates with PB1 and NS1 **(Fig. 7B)**. Overexpression of PB1 or NS1 prior to IAV infection increased both viral load and endogenous PANK4 expression, as shown by immunoblotting and immunofluorescence **(Fig. 7C −F)**. Co-expression of PANK4 with PB1 or NS1 further augmented viral replication compared with PB1 or NS1 alone **(Fig. EV 7B−C)**. Notably, PB1 or NS1 expression in the absence of infection also upregulated endogenous PANK4, as confirmed by immunoblotting, immunofluorescence, and RT–PCR **(Fig. 7G−H and Fig. EV 7D−E)**. These results identify PB1 and NS1, a viral protein induces PANK4 and support IAV replication, suggest how IAV developed multiple strategies to evade host innate immunity and establish IAV infection.

**Figure 7.**
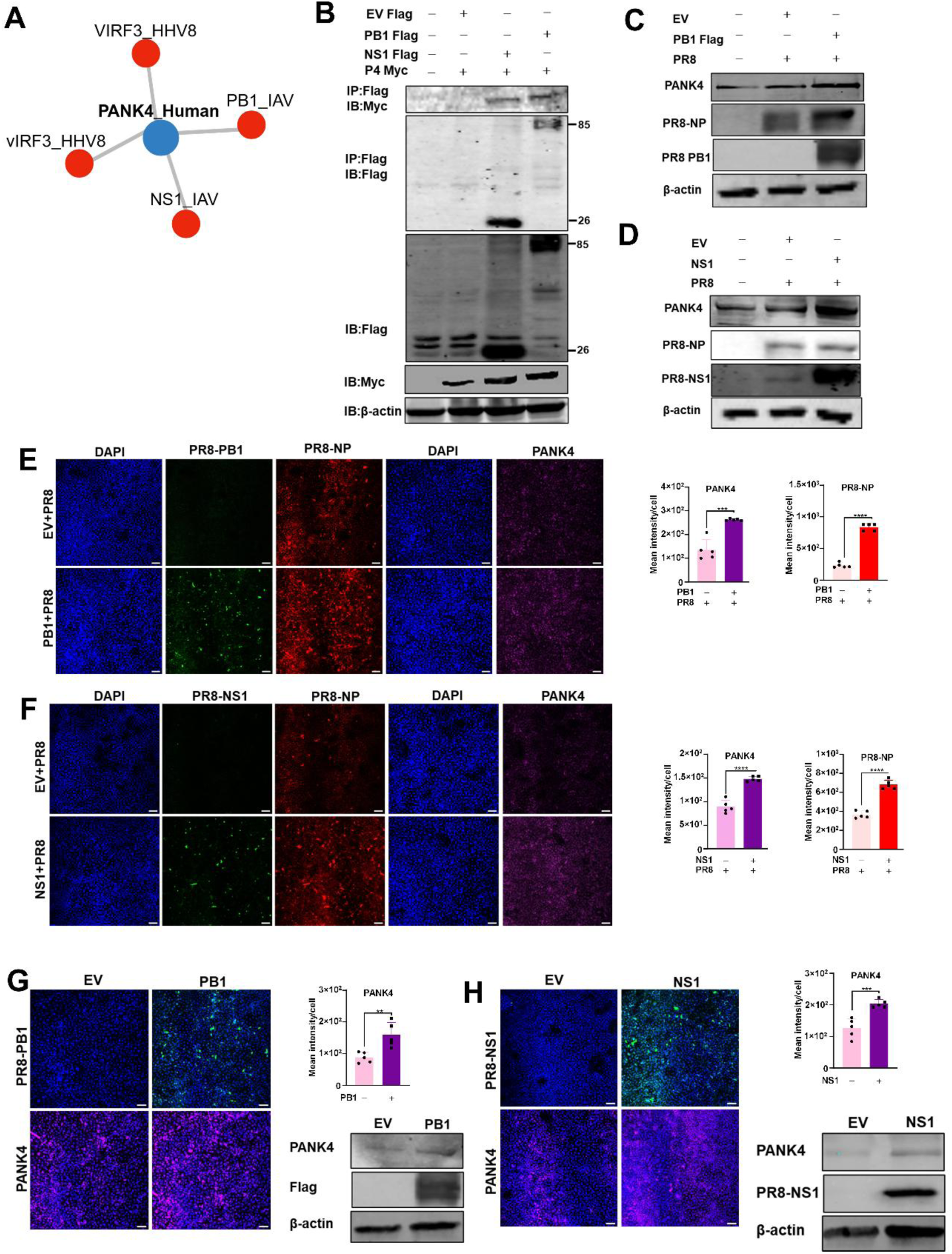
PANK4 interacts with influenza virus PB1 and NS1 to enhance viral replication. (A) In silico prediction using the Human Virus Interaction Database (HVIDB) showing potential PANK4 interactors. (B) HEK293T cells were transfected with construct encoding influenza viral proteins PB1 or NS1 (Flag tagged) or PANK4 (Myc tagged) and subjected for co-immunoprecipitation with PANK4 (Myc tagged) analysed by immunoblot (C-F) A549 cells were transfected with PB1 or NS1 overexpression plasmids and infected with PR8 (MOI 1, 24 h). PR8-NP, PB1, NS1 and endogenous PANK4 expression were analysed by immunoblot and immunofluorescence microscopy (scale bar, 100 µm) by using indicated antibodies; mean fluorescence intensity per cell quantified using QuPath-0.4.3 from five independent replicates. (G-H) A549 cells were transfected with PB1 or NS1 overexpression plasmids. PB1, NS1 and endogenous PANK4 expression were analysed by immunoblot and immunofluorescence microscopy (scale bar, 100 µm) by using indicated antibodies; mean fluorescence intensity per cell quantified using QuPath-0.4.3 from five independent replicates. Data are representative of three independent experiments.

## DISCUSSION

Innate immunity is deeply integrated with cellular metabolism, and perturbations in metabolic pathways can reshape host–pathogen interactions. Here, we identify PANK4 as a previously unrecognized regulator of antiviral innate immunity that links metabolic reprogramming to innate immune signalling through PRRs. PANK4 belongs to pantothenate kinase family, the rate-limiting enzymes of the CoA biosynthetic pathway derived from vitamin B5 (Dibble *et al*., 2022). Unlike canonical PANKs, PANK4 lacks kinase activity and acts as a metabolic pseudoenzyme. RNA viruses, particularly IAV, together with glucose or PA, induce its expression. PANK4 enhances replication by suppressing UNC93B1-dependent TLR signalling and directly engaging viral PB1 and NS1, establishing it as a metabolic–immune hub co-opted for immune evasion.

Viruses are known to remodel host metabolic networks to efficiently support their replication. Glycolysis and cofactor biosynthesis are frequently upregulated to meet the energetic and anabolic requirements for viral propagation (Mayer *et al*, 2019). Our findings extend this paradigm by identifying PANK4 as a critical mediator of IAV-induced glycolytic reprogramming. PANK4 expression was strongly induced upon infection and further amplified by PA supplementation. Functional assays revealed that PANK4 enhanced glucose uptake through the expression of glucose transporter, GLUT1 and glycolytic regulators, HK2. In contrast, PANK4 deficiency markedly impaired glucose uptake, curtailed viral replication, and suppressed glycolytic gene expression. Importantly, inhibition of glycolysis attenuated PANK4 expression itself, suggesting the existence of a feed-forward circuit linking viral infection, metabolic reprogramming, and PANK4 induction. In vivo, PA supplementation amplified this effect, leading to higher viral loads and worsened pathogenesis, implicating the Coenzyme A biosynthetic pathway as a nutrient-sensitive checkpoint of viral pathogenesis.

Beyond metabolic role, PANK4 directly regulates innate immune signalling. Mechanistically, we demonstrate that PANK4 directly interacts with UNC93B1, a trafficking chaperone required for transportation of endosomal TLR7 and TLR9. PANK4 loss resulted in increased UNC93B1 abundance, augmented trafficking, and enhanced downstream cytokine production. Conversely, PANK4 overexpression suppressed these responses, diminishing the induction of antiviral responses in terms cytokines. These results identify PANK4 as a crucial negative regulator of nucleic acid–sensing TLR signalling pathways in all cell-types tested in this study, providing a mechanism by which metabolic cues can fine-tune antiviral immunity. This finding situates PANK4 within a growing class of metabolic enzymes that exert non-canonical, functions in immune regulation.

IAV has evolved to exploit PANK4 to its own advantage. We identified the viral proteins NS1 and PB1 as binding partners of PANK4, both of which were sufficient to induce its expression and enhance viral replication. These interactions suggest that IAV co-opts PANK4 as part of its strategy to couple host metabolic remodelling along with the targeting innate immunity for viral propagation. PB1 interaction with PANK4, in turn, highlights the convergence of viral polymerase function with host metabolic remodelling, suggesting a coordinated strategy to couple replication machinery with nutrient availability. Notably, the ability of individual PANK4 domains to interact with UNC93B1 was insufficient to promote viral replication; only the full-length protein was competent, underscoring the requirement for structural integrity in orchestrating its dual metabolic and immunological functions.

Taken together, our results establish PANK4 as a central regulator at the intersection of metabolism and innate antiviral immunity. By driving glucose uptake and glycolysis, promoting viral replication, and restraining TLR7 and TLR9 mediated responses through interaction with UNC93B1, PANK4 emerges as a host factor actively subverted by influenza virus. These findings broaden the conceptual framework of immunometabolism, moving beyond metabolic flux and nutrient sensing to encompass the regulatory roles of enzymes traditionally considered catalytically inactive or regulatory in nature. The evolutionary conservation of PANK4 and its responsiveness to PA highlight the CoA biosynthetic pathway as a nutrient-sensitive axis linking metabolic state to antiviral defense.

The identification of PANK4 as both a metabolic checkpoint and immunological regulator highlights selective pressure viruses exert on host metabolic enzymes and suggests metabolic pathways are recurrent viral targets. Given its dual role in metabolism and immunity, strategies that limit PA availability, block PANK4 induction, or disrupt its interaction with viral proteins may offer new antiviral avenues. More broadly, this work shows how viral exploitation of metabolic enzymes uncovers unexpected immune-regulatory mechanisms, enhancing our understanding of the host–pathogen interface and opening opportunities for therapeutic targeting of immunometabolism.

## METHODS

### Reanalysis of publicly available data from NCBI-GEO database

High throughput sequencing data were reanalysed from the NCBI Gene Expression Omnibus (GEO) dataset GSE225643 (Li *et al*., 2023), which profiles glucose-regulated gene expression in MDA-MB-231 cells. The dataset analysed using the GEO2R web tool (NCBI) to identify differentially expressed genes under glucose-supplemented versus glucose-starved conditions. Expression values were normalized, and genes with an adjusted P value < 0.05 and |log₂ fold change| ≥ 1 were considered significantly regulated and plots were generated.

### Sequence alignments and comparisons

Protein sequences were aligned using the MultAlin multiple-sequence alignment program. The resulting alignments were visualized and annotated with ESPript 3.0, which generates publication-quality renderings (Robert & Gouet, 2014).

### Cell culture, transfection and reagents

A549 human alveolar basal epithelial cells (ATCC CCL-185), HEK293T human embryonic kidney cells (ATCC CRL-3216), HepG2 human hepatoblastoma cells (ATCC HB-8065), THP-1 human monocytes (ATCC TIB-202), and RAW264.7 murine macrophages (ATCC TIB-71) were maintained in Dulbecco’s modified Eagle’s medium (DMEM) supplemented with 10% fetal bovine serum (FBS) and 1% penicillin–streptomycin. Transfections with plasmid DNA (including shRNA), small interfering RNA (siRNA) were carried out using Lipofectamine 3000 (Invitrogen), according to the manufacturer’s protocol. For electroporation, THP-1 cells and hPBMCs (1 × 10^6^) were resuspended in Opti-MEM (Invitrogen) containing siRNA or plasmid DNA and subjected to two pulses of 1,000 V for 0.5 ms with a 5-s interval using a Gene Pulser Xcell electroporation system (Bio-Rad). Following electroporation, cells were transferred to RPMI medium supplemented with 10% FBS and cultured under standard conditions. For starvation cells were cultured in DMEM and subjected to glucose starvation using glucose-free DMEM (Thermo Fisher). Recombinant IFN-β (IF014 sigma), human TNFα, (H8916, sigma), Ecoli LPS (L2630, sigma) or CpG oligodeoxynucleotide (ODN, Medchem Agatolimod (HY-150218)) were stimulated as indicated amount. FLAG-tagged constructs of PB1, NP, NS1, and M2 were kindly provided by Shitao Li (Wang *et al*, 2017).

### Viruses and infection

A/PR8/H1N1 influenza virus (multiplicity of infection [MOI] = 1), generated as described previously(Kumar *et al*, 2022), and Newcastle disease virus expressing GFP were used in this study. NDV-GFP was kindly provided by Professor Peter Palese (Icahn School of Medicine at Mount Sinai, New York, USA). Cells were infected in serum-free Dulbecco’s modified Eagle’s medium (DMEM; Invitrogen) for 1 h, washed with phosphate-buffered saline (PBS), and subsequently cultured in DMEM supplemented with 1% fetal bovine serum (FBS; Invitrogen) and 1% penicillin–streptomycin (Invitrogen). The replacement medium for PR8 infection was supplemented with TPCK-treated trypsin (1 μg/ml) to facilitate viral entry and replication. For in vivo experiments, mice were intranasally infected with 100 plaque-forming units (PFU) of PR8 virus in 40 μL PBS under light anaesthesia(Galani *et al*, 2022). Dengue virus (strain DENV-2, New Guinea C) was used in this study. The virus was rescued from a reverse genetic system kindly provided by Prof. Andrew D. Davidson (University of Bristol)(Davidson, 2014).

### Mice

CD1 (Hylasco), C57BL/6 (NBRC, India), Il6⁻/⁻ (Jackson Laboratory), and Ifnar⁻/⁻ (Jackson Laboratory) mice were used in this study. All animal experiments were approved by the Institutional Animal Ethics Committee (IAEC) and performed in accordance with institutional guidelines. Mice were bred and maintained under specific pathogen-free conditions at the animal facility of IISER Bhopal, with ad libitum access to food and water, controlled ambient temperature (21 ± 1 °C), relative humidity (40–60 ± 5%), and a 12 h light/dark cycle. Both male and female mice (4–6 weeks old) were randomly assigned to experimental groups (n = 6 per group). To minimize sex bias, each group consisted of three male and three female mice, analysed separately but included equally in all experimental conditions.

### PA treatment

D-Pantothenic acid hemicalcium salt (Cat. no. 21210-5G-F; Sigma) was administered intraperitoneally at an initial loading dose of 500 mg/kg one day before infection, followed by a maintenance dose of 100 mg/kg per day until sample collection (Bais *et al*, 2022).

### BAL Fluid collection

Mice were euthanized at indicated time points, and the trachea was surgically exposed and cannulated with a sterile 20-gauge catheter. Bronchoalveolar lavage (BAL) was performed by instilling 1 ml of cold sterile PBS into the lungs, followed by gentle aspiration. The procedure was repeated three times and recovered lavage fluids were pooled. BAL samples were centrifuged at 500g for 10 min at 4 °C to separate cells from supernatants, which were stored at −80 °C until further analysis (Galani *et al*., 2022).

### Isolation of primary cells from mouse lung tissue

Lungs from 4–6-week-old mice were harvested under sterile conditions. After euthanasia, the pulmonary circulation was perfused via the right ventricle with cold PBS to remove blood. Lungs was minced and digested in media containing DNase I (0.01% w/v), collagenase and diaspase solution (Himedia, TCL142) (1:10 ratio) and incubated in 37°C for 30-45 min. Cell suspensions were passed through 40-µm strainers, red blood cells were removed by brief ammonium–chloride lysis, and cells were washed in PBS/2% FBS and cultured in DMEM supplemented with 10% FBS and 1% penicillin–streptomycin at 37°C (Fuentes-Mateos *et al*, 2023). The non-adherent cells were removed with PBS wash after 24 h and the adherent cells were supplemented with culture media for primary lung cells (Edelman & Redente, 2018).

### PBMC and BMDC isolation

Human PBMCs were obtained from healthy donors using EDTA-coated blood collection tubes and isolated by density gradient centrifugation with Histopaque-1077 (Sigma) according to the manufacturer’s instructions. Cells were cultured in RPMI 1640 medium supplemented with 10% FBS and infected at a MOI of 1. For BMDCs, After euthanasia, the four limbs of mice was removed, and the bone marrow was extracted by flushing PBS with 25G needle syringe. The bone marrow cells were dissociated and washed with PBS at 1000 x g for 5 min. The cells were suspended in DMEM supplemented with 10% FBS, 1% penicillin–streptomycin, 25ng/mL GM-CSF and incubated at 37°C for 6-7 days for mature BMDCs (Diotallevi *et al*, 2022).

### Quantitative RT PCR

Intracellular RNA was extracted using TRIzol reagent (Invitrogen), and RNA from culture supernatants and BAL fluid was isolated with TRIzol LS reagent (Invitrogen). cDNA was synthesized using the High-Capacity cDNA Reverse Transcription Kit (Thermo Fisher Scientific) according to the manufacturer’s instructions. Gene expression was measured by quantitative real-time PCR (RT-PCR) with gene-specific primers and PowerUp SYBR Green Master Mix (Thermo Fisher Scientific). The primers used for RT-PCR is attached in table **(Appendix Table S 3)**

### Digital PCR

RNA was extracted using TRIzol reagent (Invitrogen), and cDNA was synthesized with the High-Capacity cDNA Reverse Transcription Kit (Thermo Fisher Scientific) following the manufacturer’s instructions. Digital PCR was performed on A549 cell cDNA using PANK4-specific primers and the QIAcuity EG PCR Kit (QIAGEN, 250111).

### Enzyme-linked immunosorbent assay (ELISA)

Culture supernatants from A549 cells were collected, and levels of IP-10 and TNFα were quantified using ELISA kits (BD OptEIA™ Human IP-10 ELISA Set, 550926; BD OptEIA™ Human TNFα ELISA Set, 555212) following the manufacturer’s instructions.

### Glucose Uptake Assay

Glucose uptake was assessed using 2-Deoxy-2-[(7-nitro-2,1,3-benzoxadiazol-4-yl)amino]-D-glucose (2-NBDG, 72987, Sigma-Aldrich) and 2-Deoxy-D-glucose (2-DG, D8375, Sigma-Aldrich). A549 cells were incubated with 2-NBDG (200 μM; 20 min; 37 °C) in PBS supplemented with 1 mM MgCl₂ and 1 mM CaCl₂. After washing with PBS, intracellular uptake of 2-NBDG was analysed by FACSAria III flow cytometer (Becton Dickinson) and High-content imaging was performed using the CellInsight CX7 LZR Pro High-Content Screening (HCS) Platform (Thermo Fisher Scientific) equipped with automated fluorescence acquisition and analysis capabilities. For inhibition assays, cells were treated with 2-DG as indicated in the figure legends, and downstream effects on gene expression were evaluated by quantitative RT–PCR (Bhatt *et al*., 2022).

### Immunoprecipitation (IP) and Immunoblotting analysis

Cells were harvested 36 h post-infection, washed with PBS, and lysed in ice-cold lysis buffer supplemented with a protease inhibitor cocktail (11836145001, Roche). Lysates were cleared by centrifugation at 15,000 g for 10 min at 4 °C, and supernatants were collected. Protein concentrations were determined using the Bradford assay, and ∼30 µg of protein per sample was resolved by SDS–PAGE. Immunoblotting was performed using antibodies against PANK4 (CST, 12055), RIG-I (Thermo, MA5-31715), GFP, IAV-NP (Thermo, PA5-32242), TLR7 (CST, 5632), UNC93B1 (Thermo, PA5-20510), FLAG (Sigma, F1804), MYC (Sigma, C3956), GAPDH (Sigma, G9545), IAV-NS1 (Thermo, PA5-32243), Calreticulin (Thermo, MA5-45066) and β-actin (A1978, Sigma-Aldrich). Secondary anti-mouse and anti-rabbit IgG antibodies were obtained from LI-COR, and blots were visualized using the LI-COR imaging system. For mouse tissue samples, ∼5 mg of tissue was homogenized in a mortar and pestle and lysed using the same buffer with SDS substituted for NP-40.

For immunoprecipitation, clarified lysates were incubated with Anti-FLAG M2 magnetic beads (M8823) at 4 °C with gentle rotation. Beads were washed thoroughly, and bound proteins were eluted, denatured, and analysed by immunoblotting as described above.

### Flow cytometry

Cells were harvested 36 hours post-infection, trypsinized, and fixed with 4% para formaldehyde for 10 minutes. Fixed cells were permeabilized with 0.1% Triton X-100 for 10 minutes, blocked with 5% FBS for 40 minutes, and incubated with primary antibodies for 40 minutes followed by secondary antibodies for 30 minutes. Samples were analysed on a FACSAria III flow cytometer (Becton Dickinson), and data were processed using FlowJo software.

### Confocal and HCS microscopy

Cells were seeded on coverslips (confocal) or in 96-well plates (HCS), fixed in 4% paraformaldehyde for 15 min, permeabilized with 0.1% Triton X-100 and blocked in 5% BSA. Samples were incubated with primary antibodies and appropriate fluorophore-conjugated secondary antibodies, and nuclei were counterstained with DAPI.Confocal imaging was performed using a Zeiss LSM confocal microscope with a 100× oil-immersion objective for high-resolution colocalization analysis. Colocalization was quantified in ImageJ/Fiji (e.g., Pearson’s correlation coefficient) on background-subtracted images. HCS imaging was performed using a 10× objective to quantify viral load and overall phenotypes across wells. For HCS, multiple fields per well were acquired and analysed with fiji/imagej software and figure 6 and 7 10x images were quantified using QuPath-0.4.3 software.

### Statistical analysis

All experiments were performed with appropriate controls or mock-transfected samples and independently repeated two to three times. Statistical analyses were carried out using GraphPad Prism software, version 8. Comparisons between two groups were performed with an unpaired two-tailed Student’s t-test, while comparisons among three or more groups were evaluated by ANOVA. A P value < 0.05 was considered statistically significant. Statistical significance in figures is denoted as follows: *P < 0.05, **P < 0.01, ***P < 0.001; ns, not significant.

## Acknowledgements

We thank R. Fouchier for providing the A/PR8/H1N1 reverse genetics system and P. Palese for generously sharing the GFP-expressing Newcastle disease virus (NDV-GFP). We also thank Shitao Li for providing constructs expressing IAV genes. We acknowledge the Central Instrumentation Facility at IISER Bhopal for technical support. We are grateful to Dr. Anamika Mishra and Dr. Ashwin A. Raut (ICAR-NIHSAD) for their assistance with virus propagation. We also thank Pandikannan K. and Madhavan P. for their help with the initial analysis. This work was partially supported by the Indian Council of Medical Research (ICMR; Grant No. ICMR/BIO/2023-2024/98) and by institute funding from IISER Bhopal. R.C. acknowledges support from the University Grants Commission (UGC), and A.M. from the Department of Biotechnology (DBT), Government of India.

## Author contributions

**Riya Chaudhary**: Conceptualization; Methodology; Formal analysis; Investigation; Writing— original draft; Writing—review and editing.

**Aparna Meher**: Conceptualization; Methodology; Formal analysis; Investigation; Writing— review and editing.

**Pratik Katekar**: Investigation; Writing—review and editing.

**Drishiya Vats**: Animal experiment.

**Debasis Nayak**: Animal experiment.

**Himanshu Kumar**: Conceptualization; Methodology; Formal analysis; Supervision; Funding acquisition; Writing—original draft; Writing—review and editing.

## Ethics declarations

All animal experiments were performed in compliance with institutional guidelines and approved by the Institutional Animal Ethics Committee (IAEC) of IISER Bhopal (Approval No. 2025-IISERB-7.10-IAEC). All experiments involving influenza virus and Newcastle disease virus were conducted under biosafety guidelines with approval from the Institutional Biosafety Committee (IBSC) of IISER Bhopal (Approval No. IBSC/IISERB/2025/Meeting-1/05).

## Figure Expanded View

**Figure EV 1.**
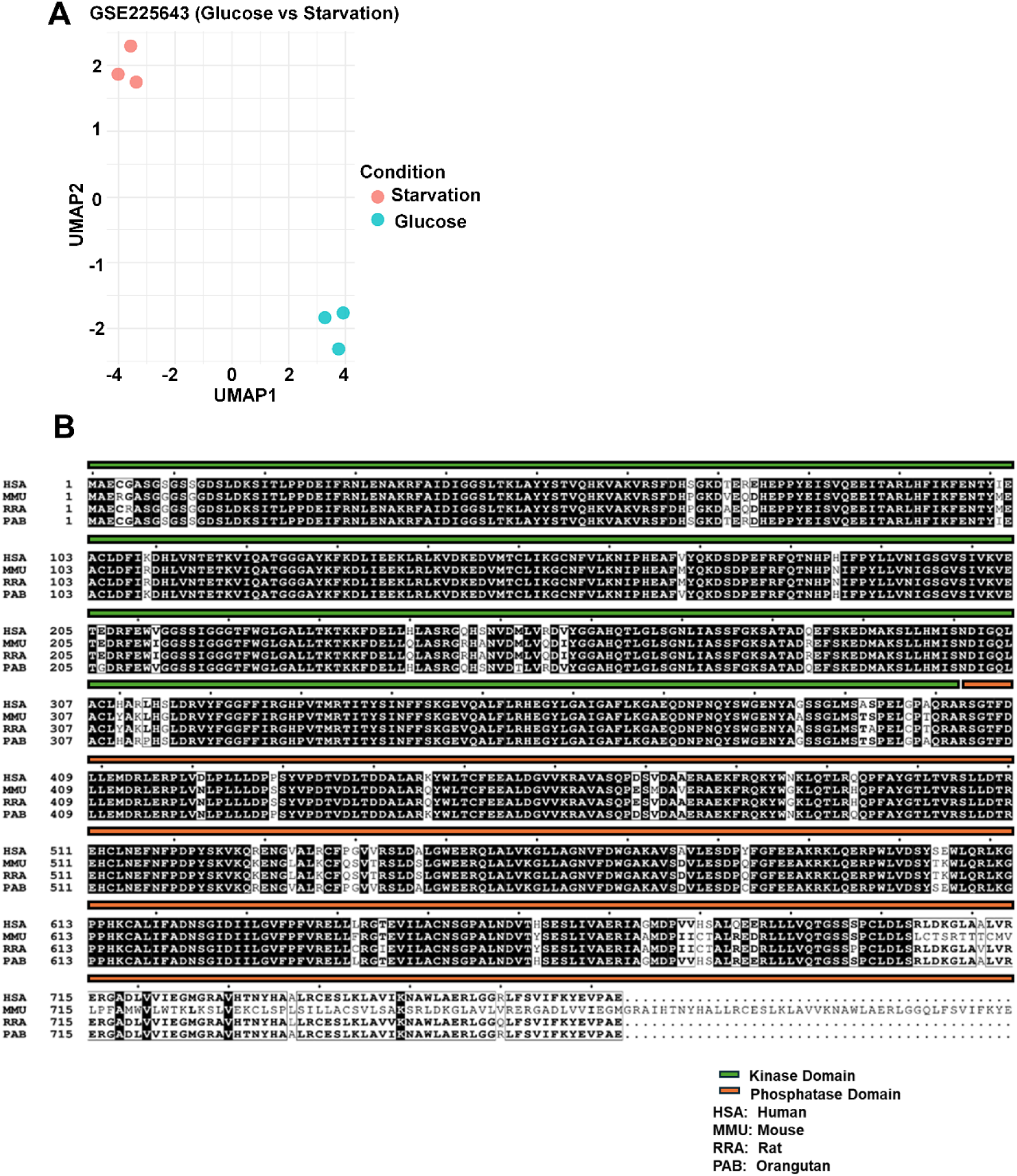
Identification and evolutionary conservation of PANK4. (A) Uniform manifold approximation and projection (UMAP) plot showing sample clustering from the glucose-regulated gene expression dataset (GSE225643). Samples correspond to glucose-supplemented (blue) and glucose-starved (red) conditions. (B) Multiple sequence alignment was performed using ESPript 3.0, highlighting the high conservation of PANK4 among mammalian species including Homo sapiens (HSA), Mus musculus (MMU), Rattus norvegicus (RRA), and Orangutan (PAB). Fully conserved residues are shaded in black, while similar residues are boxed. Functionally important domains are marked in green and orange, respectively. Data are presented as the mean ± SEM from triplicate samples of a single analysis. *** p<0.001 by two-tailed unpaired Student’s t-tests.

**Figure EV 2.**
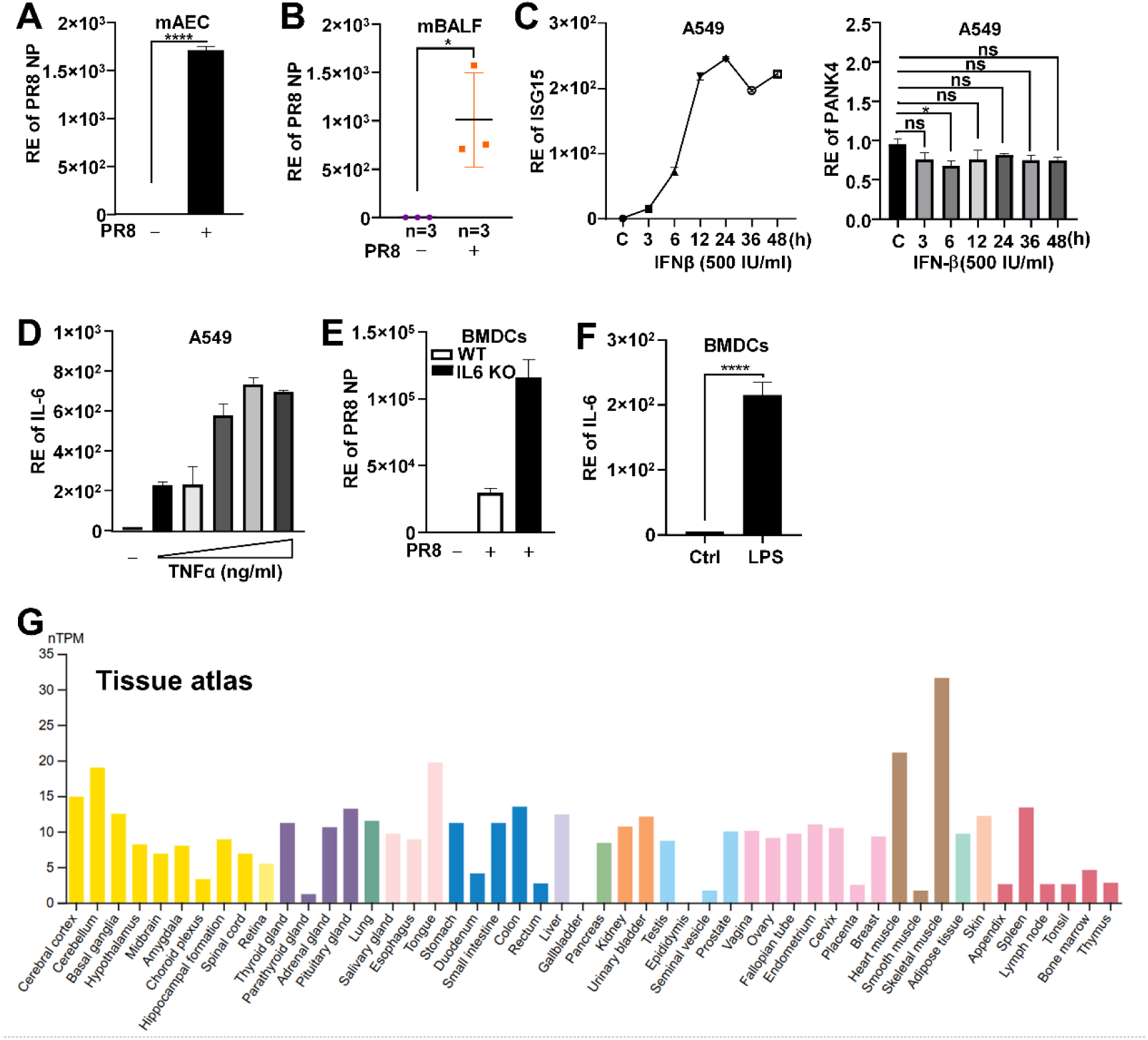
PANK4 induction is independent of type I IFN and inflammatory cytokines. Ex vivo, mAECs (A) were isolated and infected with PR8 (MOI 1) for 24 h. (B) In vivo, PR8 infected BAL fluid (PFU 100, dpi 3) was collected, after which the RNA level of PR8-NP were measured. (C) A549 cells were treated with IFN-β (500 IU/ml) for indicated timepoint and ISG15 (Positive control) or RNA level of PANK4 were analysed. (D) A549 cells stimulated with TNF-α (10–100 ng/ml) for 12 h and analysed for IL-6 transcript level. (E) WT and IL6^-/-^ BMDCs with or without PR8 (MOI 1) infection, post infection RNA level of PR8-NP were measured. (F) BMDCs stimulated with LPS (100 ng/ml) for 24 h and IL-6 expression were measured. (G) PANK4 expression across human tissues based on publicly available Tissue Atlas profiling. A-F were analysed by RT-PCR. Data are presented as the mean ± SEM from triplicate samples of a single experiment and representative of three to five independent experiments. **** p<0.0001, * p<0.05, ns is non-significant by two-tailed unpaired Student’s t-tests.

**Figure EV 3.**
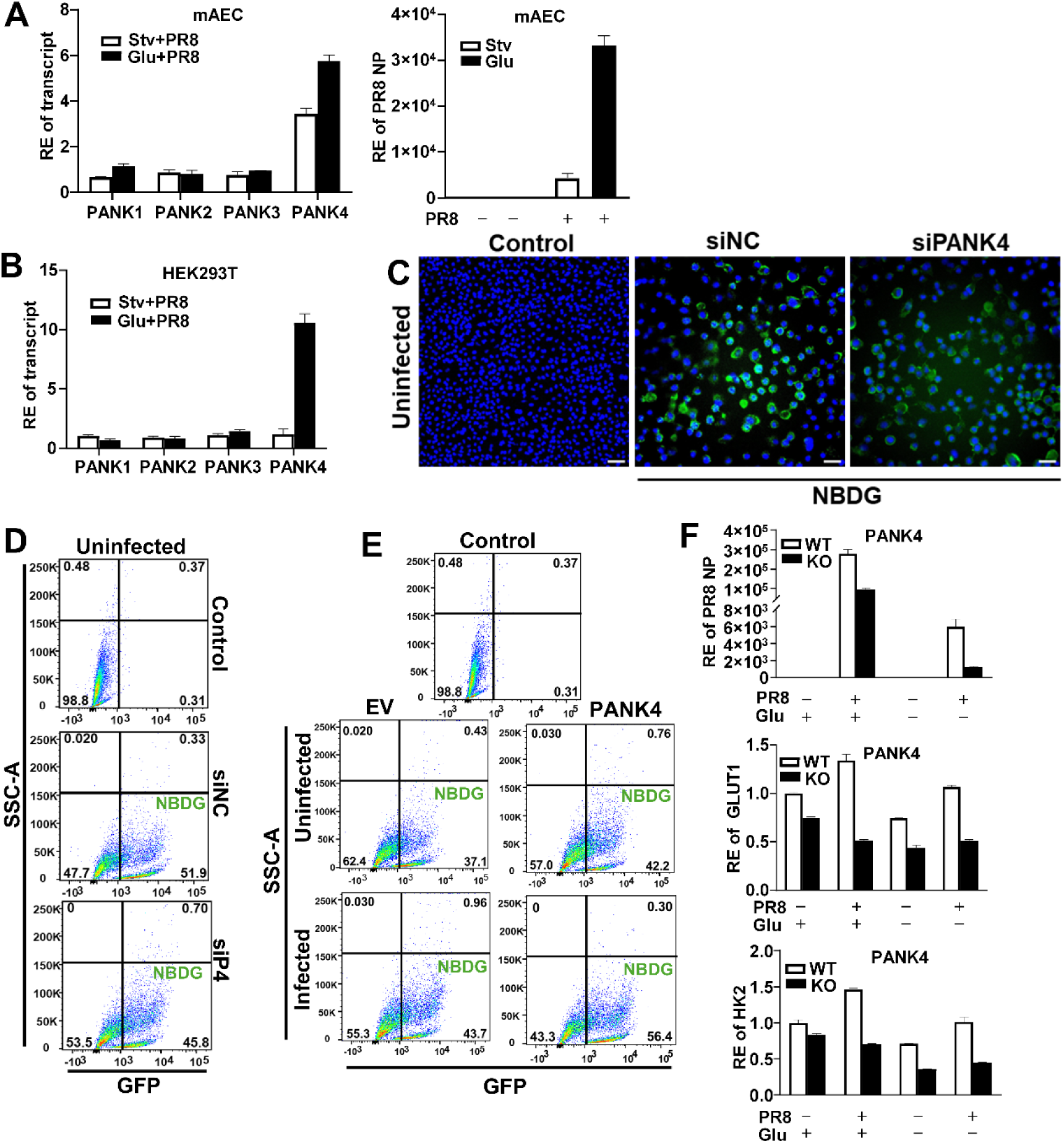
PANK4 uniquely responds to glucose availability and viral infection. Ex vivo mAECs (A) and HEK293T (B) cells were cultured under glucose-supplemented versus glucose-starved (12 h) conditions and infected with PR8 (MOI 1, 12 h). RNA level of PANK isoforms and PR8-NP were analysed. A549 cells were transfected with siPANK4 (30 nM) and subjected to starvation (6 h) and Glucose uptake (2-NBDG) were measured by immunofluorescence microscopy (C) and flow cytometry (D). (E) A549 cells were transfected with construct encoding PANK4 or infected with PR8 (MOI 1, 12 h) and subjected to starvation (6 h). Glucose uptake (2-NBDG) were measured by flow cytometry. Plots show 2-NBDG fluorescence with quadrants representing NBDG populations; numbers denote percentage of cells. (F) WT and PANK4⁻/⁻ cells comparing glucose-supplemented and glucose-starved (12 h) with PR8 infection (MOI 1, 12 h) or without infection, post infection RNA level of PR8-NP, GLUT1, and HK2 were analysed. A-B and F were analysed by RT-PCR. Data are presented as the mean ± SEM from triplicate samples of a single experiment and representative of three to five independent experiments. *** p<0.001 by two-tailed unpaired Student’s t-tests.

**Figure EV 4.**
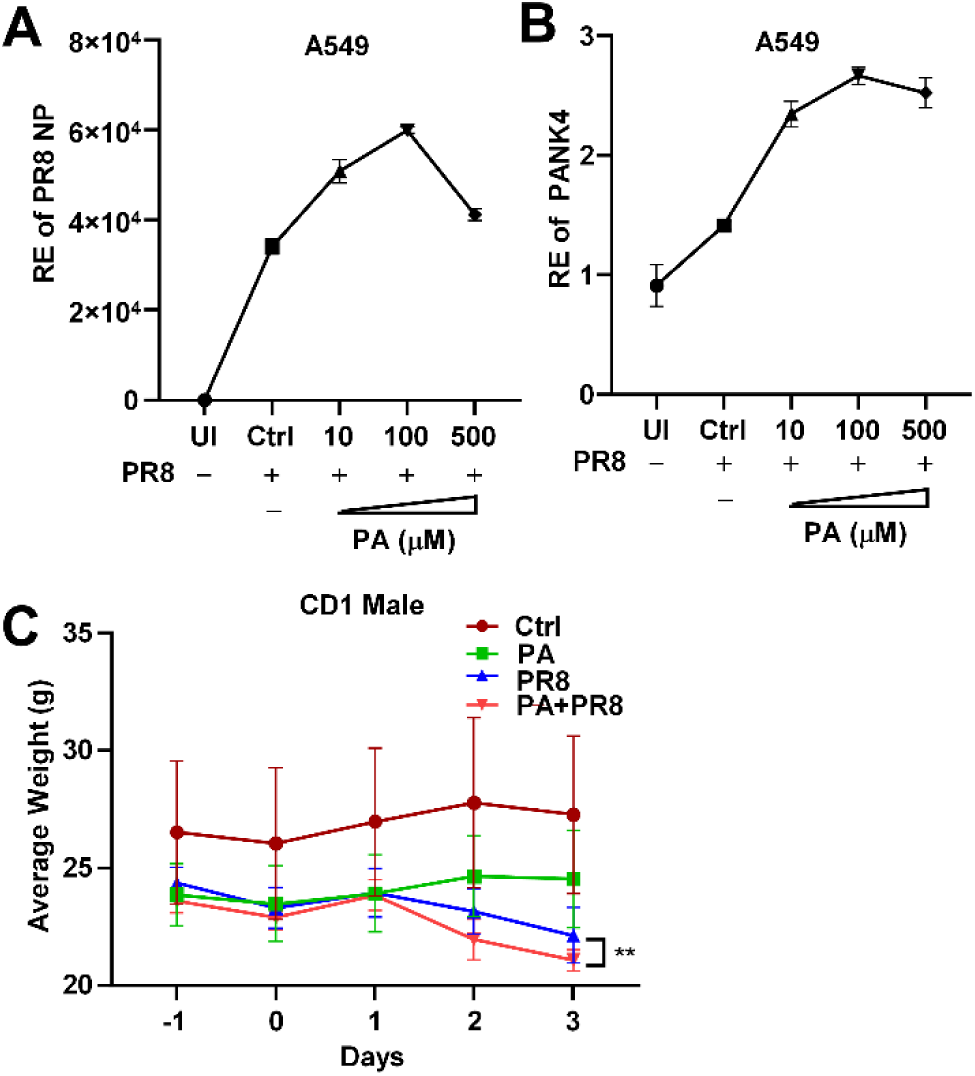
Dose- and infection-dependent induction of PANK4 by PA. (A-B) A549 cells were treated with increasing concentrations of PA for 24 h and infected with PR8 (MOI 1). RNA level of PR8-NP and PANK4 were analysed by RT-PCR. (C) Male mice body weight changes measured over the indicated time course. Data are presented with statistical significance of ** p<0.01 by two-way repeated-measures ANOVA test.

**Figure EV 5.**
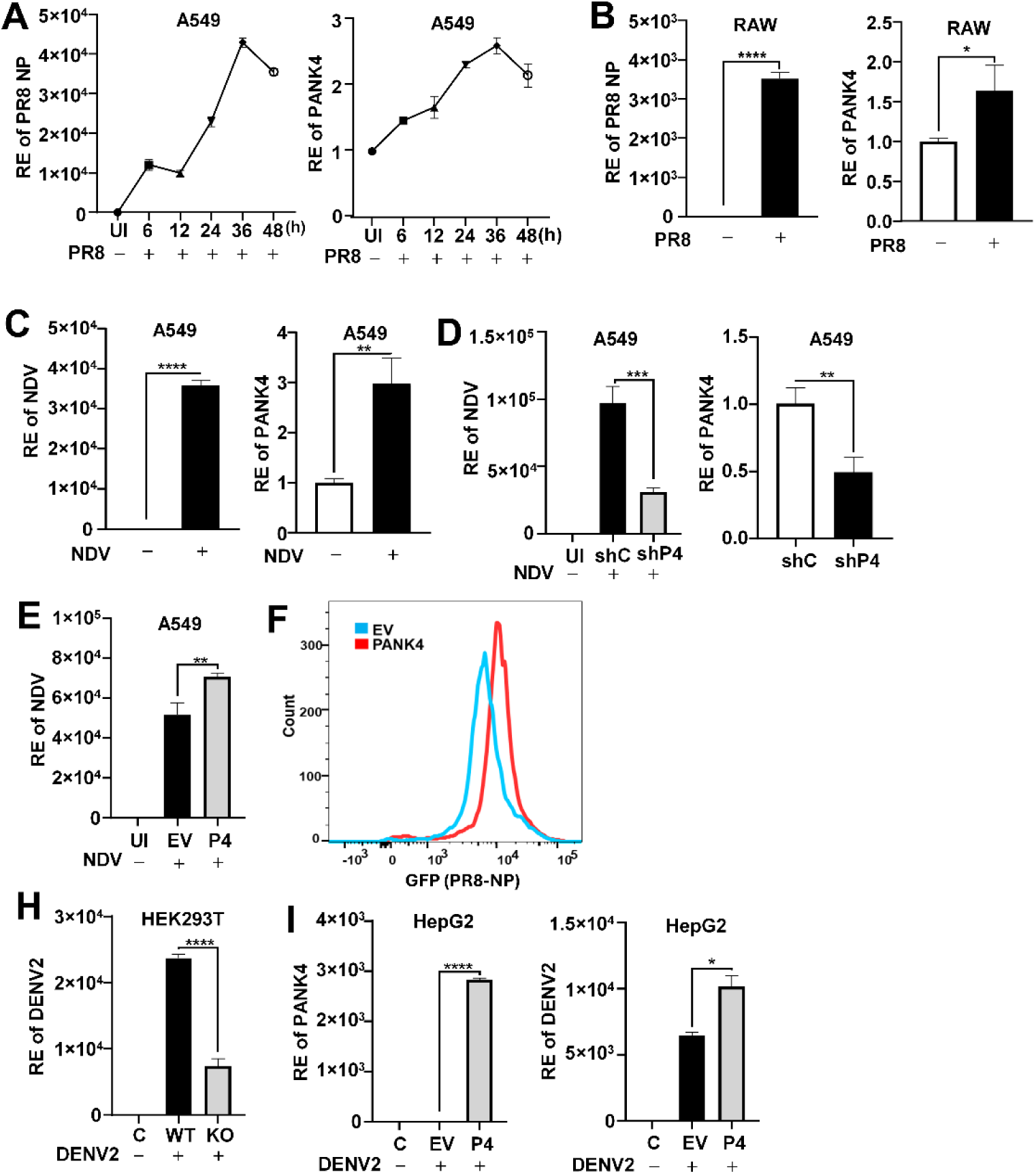
PANK4 supports viral replication across diverse RNA viruses. (A) A549 cells were infected with PR8 (MOI 1) over indicated time-course and RNA level of PANK4 and PR8-NP were analysed. RAW cells were infected with PR8 (MOI 1, 24 h) (B) and A549 cells were infected with NDV (MOI 0.5, 24 h) (C) and RNA level of PR8-NP were analysed. (D) A549 cells were transfected with shRNA of PANK4 and infected with NDV-GFP (MOI 0.5, 24 h) and NDV RNA level were analysed. (E) A549 cells were transfected with construct encoding PANK4 and infected with NDV (MOI 0.5, 24 h), post infection NDV RNA level were analysed. (F) A549 cells were transfected with construct encoding PANK4 and infected with PR8 (MOI 1, 24 h) PR8-NP expression analysed by flow cytometry. (H) PANK4⁻/⁻ cells were infected with DENV2 (MOI 5, 24 h), post infection DENV2 viral load were analysed. (I) HEPG2 cells were transfected with construct encoding PANK4 and infected with DENV2 (MOI 5, 24 h) and DENV2 viral were measured. A-E and H-I were analysed by RT-PCR. Data are presented as the mean ± SEM from triplicate samples of a single experiment and representative of three to five independent experiments. **** p<0.0001, *** p<0.001, ** p<0.01, * p<0.05 by two-tailed unpaired Student’s t-tests.

**Figure EV 6.**
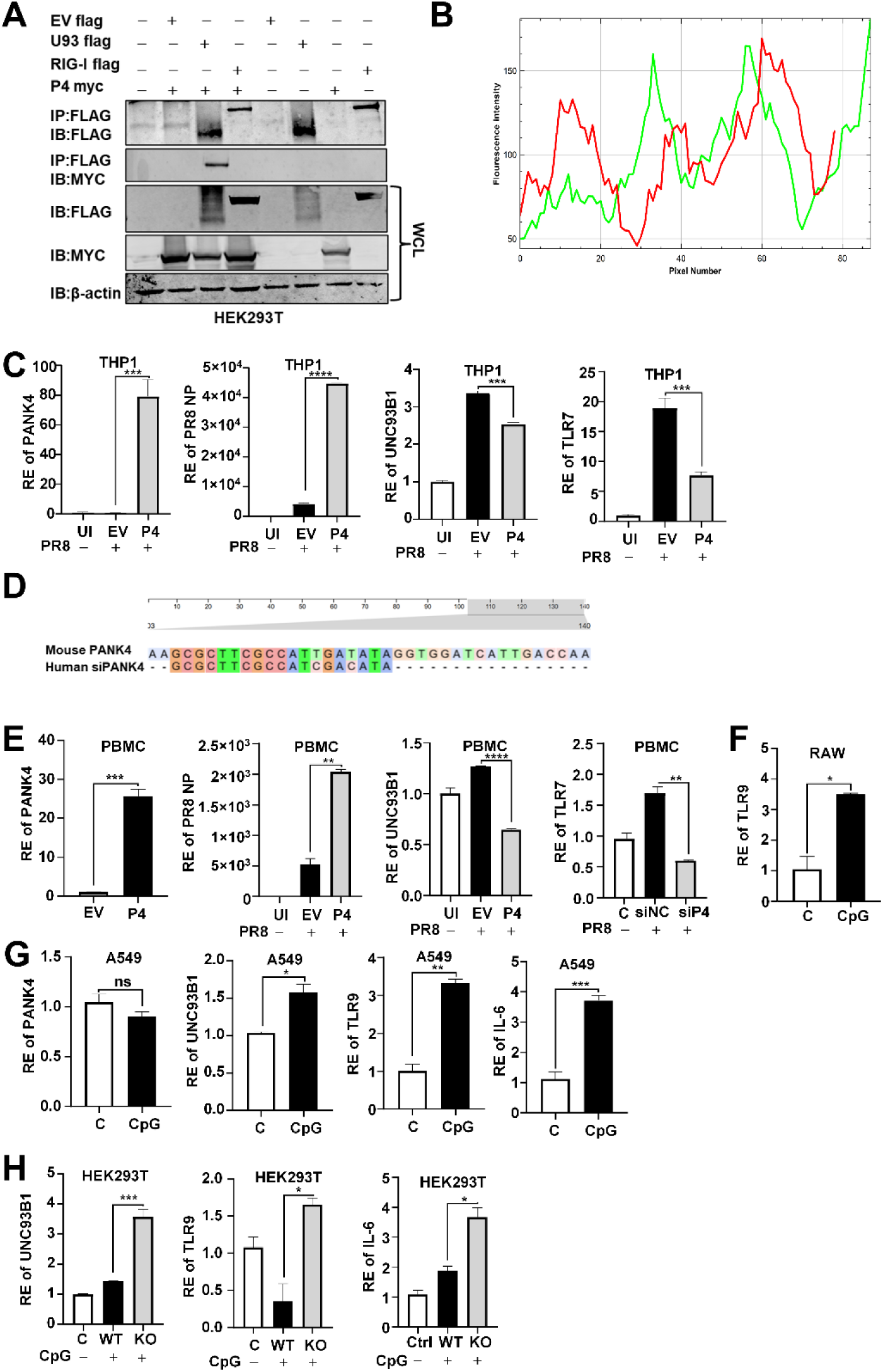
Validation of PANK4–UNC93B1 interaction and regulation of TLR pathways. (A) HEK293T cells were transfected with FLAG-tagged UNC93B1 and FLAG-tagged RIG-I MYC-tagged PANK4, and co-immunoprecipitation assays were performed. (B) Line-scan analysis of colocalization. Fluorescence intensity profiles of red and green channels along the selected line show overlapping peaks, indicating signal colocalization performed using ImageJ. (C) THP1 cells were transfected with construct encoding PANK4 and infected with PR8 (MOI 1, 24 h) and RNA level of PR8-NP, UNC93B1 and TLR7 were analysed (D) Sequence analysis demonstrated conservation of the PANK4 siRNA target site between human and mouse. (E) PBMCs were electroporated with PANK4 overexpression plasmid and infected with PR8 (MOI 1, 24 h) and RNA level of PANK4, PR8-NP, UNC93B1 and TLR7 were analysed. (F) RAW cells were stimulated with CpG ODN (5 µg/mL, 24 h) TLR9 expression were analysed. (G) A549 cells were stimulated with CpG ODN (5 µg/mL, 24 h) RNA level of PANK4, UNC93B1, TLR9, and IL-6 were analysed. (H) PANK4⁻/⁻ cells were stimulated with CpG ODN (5 µg/mL, 24 h) and RNA level of UNC93B1, TLR9, and IL-6 were analysed. C and E-H were analysed by RT-PCR. Data are presented as the mean ± SEM from triplicate samples of a single experiment and representative of three to five independent experiments. **** p<0.0001, *** p<0.001, ** p<0.01, * p<0.05, ns is non-significant by two-tailed unpaired Student’s t-tests.

**Figure EV 7.**
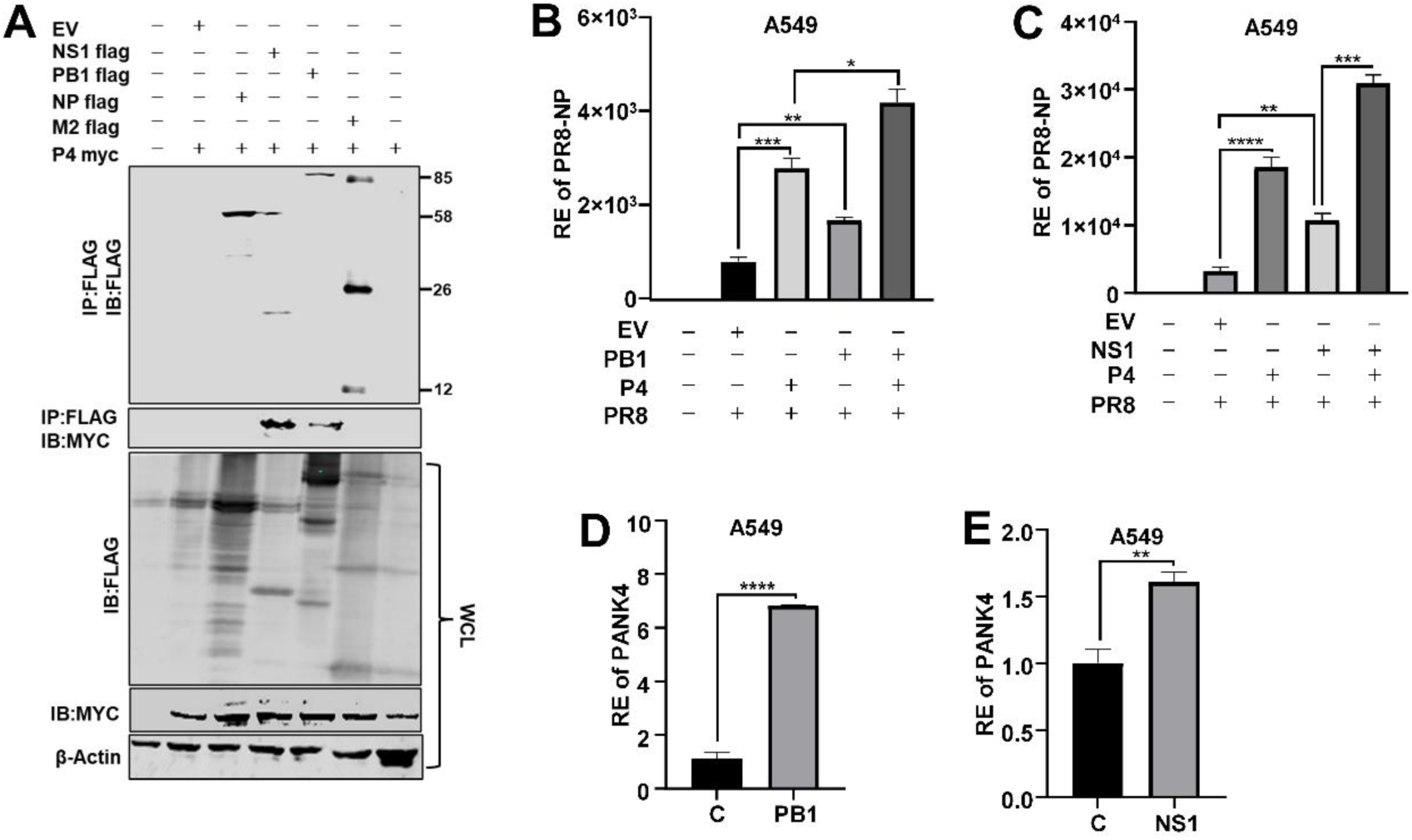
Validation of PANK4 interactions with influenza viral proteins PB1 and NS1. (A) A549 cells were transfected with flag tagged constructs encoding influenza viral proteins PB1, NP, M2, NS1 or PANK4 (Myc tagged) and subjected for co-immunoprecipitation with PANK4 (Myc tagged) analysed by immunoblot. A549 cells were transfected with flag tagged constructs encoding influenza viral proteins PB1 (B) or NS1 (C) and infected with PR8 (MOI 1, 24 h), post infection transcriptomic level of endogenous PANK4 were measured. A549 cells were transfected with flag tagged constructs encoding PB1 (D) or NS1 (E) and transcriptomic endogenous level of PANK4 were measured. B-E were analysed by RT-PCR. Data are presented as the mean ± SEM from triplicate samples of a single experiment and representative of three to five independent experiments. **** p<0.0001, *** p<0.001, ** p<0.01, * p<0.05 by two-tailed unpaired Student’s t-tests.

**Appendix Table S 1.** Host factors identified by genome-wide and kinase screens of IAV with glucose-dependent gene expression dataset (GSE225643) by GEO2R analysis.

| S. No. | Host factors | References |
| --- | --- | --- |
| 1. | CDC42BPB | (27) |
| 2. | DLG5 | (28) |
| 3. | EPHB2 | (29) |
| 4. | IL17RA | (30) |
| 5. | KPNB1 | (31) |
| 6. | MAPK1 | (32) |
| 7. | PANK4 |  |
| 8. | PHF2 | (33) |
| 9. | PIK3R4 | (34) |
| 10 | PRKACA | (35) |
| 11. | PRKCD | (36) |
| 12. | RACGAP1 | (37) |

**Appendix Table S 2.** Body weights of CD1 mice used in the study.

| **GROUP** | **Gender** | **Day -1** | **Day 0** | **Day 1** | **Day2** | **Day3** |
| --- | --- | --- | --- | --- | --- | --- |
| **Control** | Male | 26.5 | 25.56 | 27.09 | 27.8 | 27.37 |
|  | Male | 23.49 | 23.1 | 23.77 | 24.15 | 23.87 |
|  | Male | 29.6 | 29.5 | 30.08 | 31.4 | 30.6 |
|  | Female | 19.8 | 19.54 | 20.49 | 21.15 | 20.14 |
|  | Female | 23.9 | 23.22 | 22.86 | 23.36 | 22.94 |
|  | Female | 21.3 | 21.15 | 21.52 | 21.67 | 21.31 |
| **PA** | Male | 23.5 | 23.02 | 23.42 | 24.17 | 23.79 |
|  | Male | 22.78 | 22.15 | 22.59 | 23.23 | 22.94 |
|  | Male | 25.32 | 25.29 | 25.76 | 26.57 | 26.89 |
|  | Female | 19.4 | 18.6 | 19.25 | 19.46 | 18.56 |
|  | Female | 19.2 | 18.21 | 18.71 | 18.83 | 18.39 |
|  | Female | 17.82 | 17.6 | 17.9 | 18.33 | 18.43 |
| **PR8** | Male | 24.89 | 22.56 | 23.44 | 23.7 | 20.96 |
|  | Male | 23.6 | 23.11 | 23.27 | 22.04 | 22.11 |
|  | Male | 24.6 | 24.26 | 25.15 | 23.72 | 23.34 |
|  | Female | 19.4 | 18.65 | 19.36 | 18.19 | 18.39 |
|  | Female | 19.06 | 18.15 | 18.48 | 17.4 | 17.09 |
|  | Female | 19.85 | 19.72 | 20.06 | 18.6 | 17.33 |
| **PA+PR8** | Male | 23.15 | 22.52 | 23.16 | 21.06 | 21.01 |
|  | Male | 23.52 | 22.65 | 23.93 | 22.08 | 20.66 |
|  | Male | 24.12 | 23.5 | 24.46 | 22.76 | 21.56 |
|  | Female | 18.24 | 17.56 | 17.78 | 16.43 | 16.88 |
|  | Female | 18.7 | 18.32 | 18.36 | 17.18 | 16.37 |
|  | Female | 17.23 | 16.98 | 17.11 | 16.19 | 15.58 |

**Appendix Table S 3.** PANK4 interactors compiled from publicly available databases BioGRID and InnateDB.

| S. No. | PANK4 Interactors |
| --- | --- |
| 1. | EFGR 1069 |
| 2. | MAP2K2 |
| 3. | CEBPB 1070 |
| 4. | ICAM1 |
| 5. | VDR 1071 |
| 6. | UNC93B1 |
| 7. | TRIM66 1072 |
| 8. | PARK2 |
| 9. | TRIM67 1073 |
| 10 | AQP3 |
| 11. | FPR1 1074 |
| 12. | HNRNPL |
| 13. | LY86 1075 |
| 14. | PTPN6 |

**Appendix Table S 4.**
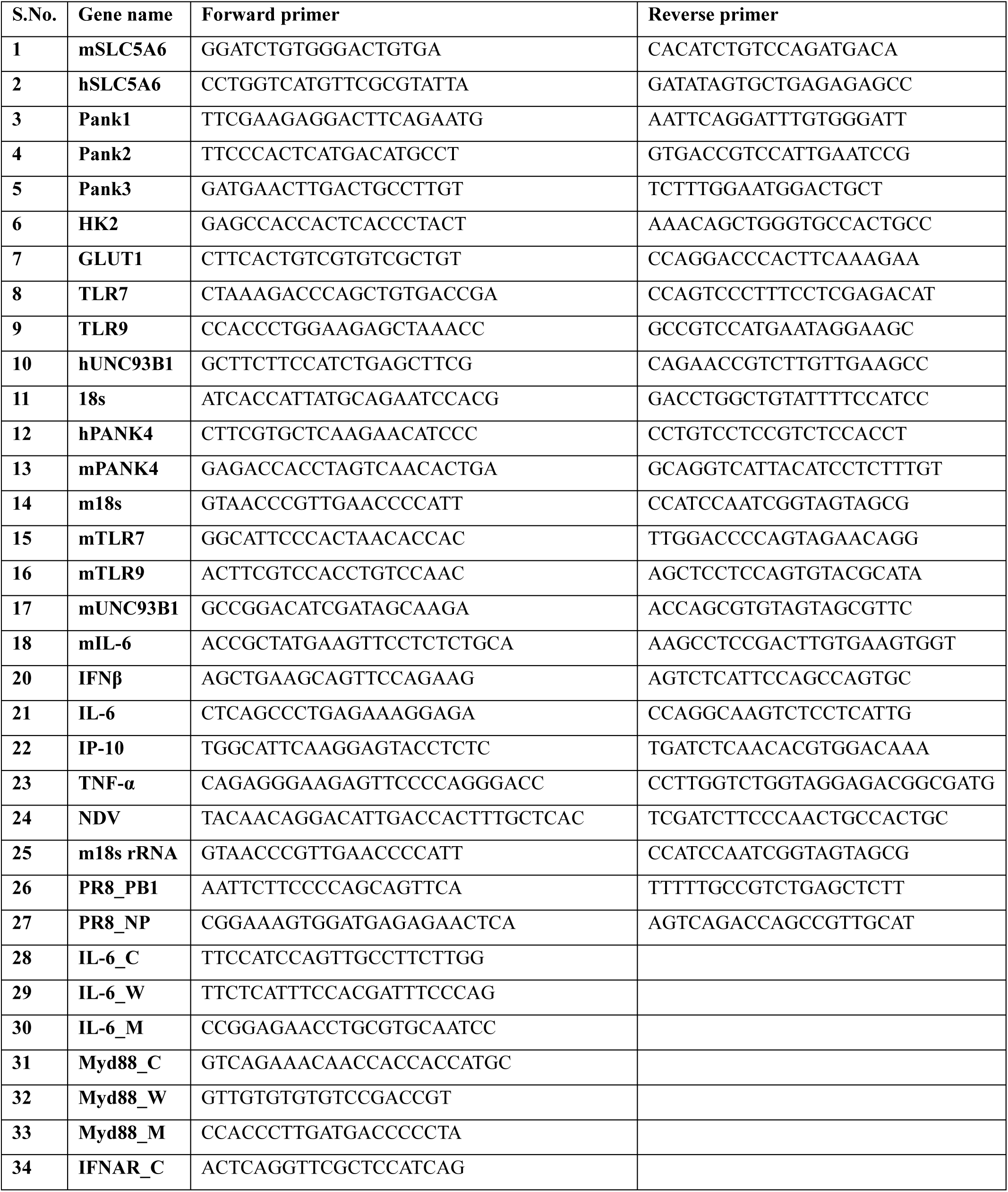
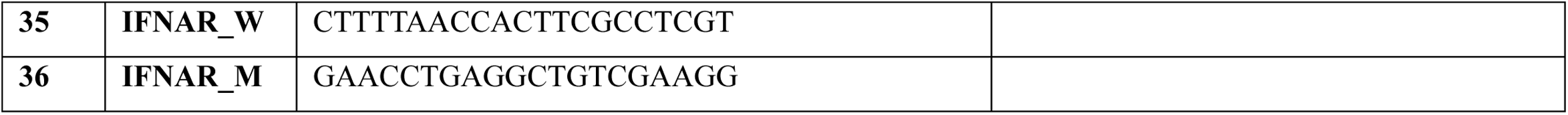
RT Primers used in this study.

